# Sequence signatures of two IGHV3-53/3-66 public clonotypes to SARS-CoV-2 receptor binding domain

**DOI:** 10.1101/2021.01.26.428356

**Authors:** Timothy J.C. Tan, Meng Yuan, Kaylee Kuzelka, Gilberto C. Padron, Jacob R. Beal, Xin Chen, Yiquan Wang, Joel Rivera-Cardona, Xueyong Zhu, Beth M. Stadtmueller, Christopher B. Brooke, Ian A. Wilson, Nicholas C. Wu

**Affiliations:** Center for Biophysics and Quantitative Biology, University of Illinois at Urbana-Champaign, Urbana, IL 61801, USA; Department of Integrative Structural and Computational Biology, The Scripps Research Institute, La Jolla, CA 92037, USA; Department of Biochemistry, University of Illinois at Urbana-Champaign, Urbana, IL 61801, USA; Department of Microbiology, University of Illinois at Urbana-Champaign, Urbana, IL 61801, USA; Carl R. Woese Institute for Genomic Biology, University of Illinois at Urbana-Champaign, Urbana, IL 61801, USA; The Skaggs Institute for Chemical Biology, The Scripps Research Institute, La Jolla, CA 92037, USA; IAVI Neutralizing Antibody Center, The Scripps Research Institute, La Jolla, CA 92037, USA; Consortium for HIV/AIDS Vaccine Development (CHAVD), The Scripps Research Institute, La Jolla, CA 92037, USA

## Abstract

Since the COVID-19 pandemic onset, the antibody response to SARS-CoV-2 has been extensively characterized. Antibodies to the receptor binding domain (RBD) on the spike protein are frequently encoded by IGHV3-53/3-66 with a short CDR H3. Germline-encoded sequence motifs in CDRs H1 and H2 play a major role, but whether any common motifs are present in CDR H3, which is often critical for binding specificity, have not been elucidated. Here, we identify two public clonotypes of IGHV3-53/3-66 RBD antibodies with a 9-residue CDR H3 that pair with different light chains. Distinct sequence motifs on CDR H3 are present in the two public clonotypes that appear to be related to differential light chain pairing. Additionally, we show that Y58F is a common somatic hypermutation that results in increased binding affinity of IGHV3-53/3-66 RBD antibodies with a short CDR H3. Overall, our results advance fundamental understanding of the antibody response to SARS-CoV-2.

## Introduction

Severe acute respiratory syndrome coronavirus-2 (SARS-CoV-2) is the etiological agent of coronavirus disease 2019 (COVID-19)^1,2^, which primarily results in respiratory distress, cardiac failure, and renal injury in the most severe cases^3,4^. The virion is decorated with the spike (S) glycoprotein, which contains a receptor-binding domain (RBD) that mediates virus entry by binding to angiotensin-converting enzyme-2 (ACE-2) receptor on the surface of host cells^1,5-7^. To mitigate the devastating social and economic consequences of the pandemic, vaccines and post-exposure prophylaxes including antibody cocktails that exploit reactivity to the S protein are being developed at an unprecedented rate. Several vaccines are currently in various stages of clinical trials^8,9^. Most notable are the mRNA vaccines from Pfizer-BioNTech and Moderna, which have been issued emergency use authorization by the Food and Drug Administration for distribution in the United States^10–12^ and the Oxford-AstraZeneca chimpanzee adenovirus vectored DNA vaccine in the United Kingdom^13–15^. In humans, most neutralizing antibodies to SARS-CoV-2 target the immunodominant RBD on the S protein^16,17^, and can abrogate virus attachment and entry into host cells^18,19^. In the past year, many RBD antibodies have been isolated and characterized from convalescent SARS-CoV-2 patients ^20–40^.

Antibody diversity is generated through V(D)J recombination^41–43^. Three genes, one from each of the variable (V), diversity (D) and joining (J) loci, are combined to form the coding region for the heavy chain. In humans, genes encoding for the V, D and J regions are denoted as IGHV, IGHD and IGHJ, respectively. Two complementarity-determining regions on the heavy chain (CDRs H1 and H2) are encoded by the V gene while the third (CDR H3) is encoded by the V(D)J junction. A similar process occurs in assembly of the coding region for the light chain except that the D gene is absent. The light chain genes also encode kappa and lambda chains that are denoted as IGKV and IGKJ, as well as IGLV and IGLJ, respectively. To further improve affinity of antibodies to an antigen, affinity maturation occurs via somatic hypermutation (SHM)^44,45^. V(D)J recombination and SHM therefore ensure a diverse repertoire of antibodies is available for an immune response to the enormous number and variety of potential antigens.

Notwithstanding this antibody diversity, some RBD antibodies with strikingly similar sequences have been found in multiple convalescent SARS-CoV-2 patients^32,46,47^. These antibodies can be classified as public clonotypes if they share the same IGHV gene with similar CDR H3 sequences^48–52^. Over the past decade, public clonotypes to human immunodeficiency virus^48^, malaria^52^, influenza^49^, and dengue virus^53^ have been discovered. Antibodies to SARS-CoV-2 RBD frequently use IGHV3-53 and IGHV3-66^23,31,47,54^, which only differ by one amino acid (i.e. I12 in IGHV3-53 and V12 in IGHV3-66). IGHV3-53/3-66 antibodies carry germline-encoded features that are critical for RBD binding – an NY motif in CDR H1 and an SGGS motif in CDR H2^31,47,54^. Nevertheless, IGHV3-53/3-66 RBD antibodies have varying lengths of CDR H3 with diverse sequences, which seem to deviate from the canonical definition of a public clonotype.

By categorizing IGHV3-53/3-66 RBD antibodies based on CDR H3 length and light chain usage, we now report on two public clonotypes of IGHV3-53/3-66 RBD antibodies, both of which have a CDR H3 length of 9 amino acids but with distinct sequence motifs. Our structural and biochemical analyses reveal that these sequence motifs on CDR H3 are associated with light chain pairing preference. We also identify Y58F as a signature SHM among IGHV3-53/3-66 RBD antibodies that have a CDR H3 length of less than 15 amino acids (Kabat numbering). As the COVID-19 pandemic continues, knowledge of public antibodies against SARS-CoV-2 can inform on therapeutic development as well as vaccine assessment.

## Results

### Two public clonotypes of IGHV3-53/3-66 RBD antibodies

In this study, we define clonotypic IGHV3-53/3-66 RBD antibodies as antibodies that share the same IGL(K)V genes and with identical CDR H3 length. Literature mining of 214 published IGHV3-53/3-66 RBD antibodies obtained from convalescent patients (Supplementary Table 1) revealed that the two most common clonotypes have a CDR H3 length of 9 amino acids and are paired with light chains IGKV1-9 (clonotype 1) and IGKV3-20 (clonotype 2), respectively (Figure 1a). Antibodies from clonotype 1 have been observed across 10 studies^22-24,32-36,40^, whereas antibodies from clonotype 2 are found across seven studies^22,24,32-34,37,40^. Interestingly, sequence logos revealed distinct sequence features of CDR H3 between clonotype 1 and clonotype 2 antibodies (Figure 1b).

**Figure 1.**
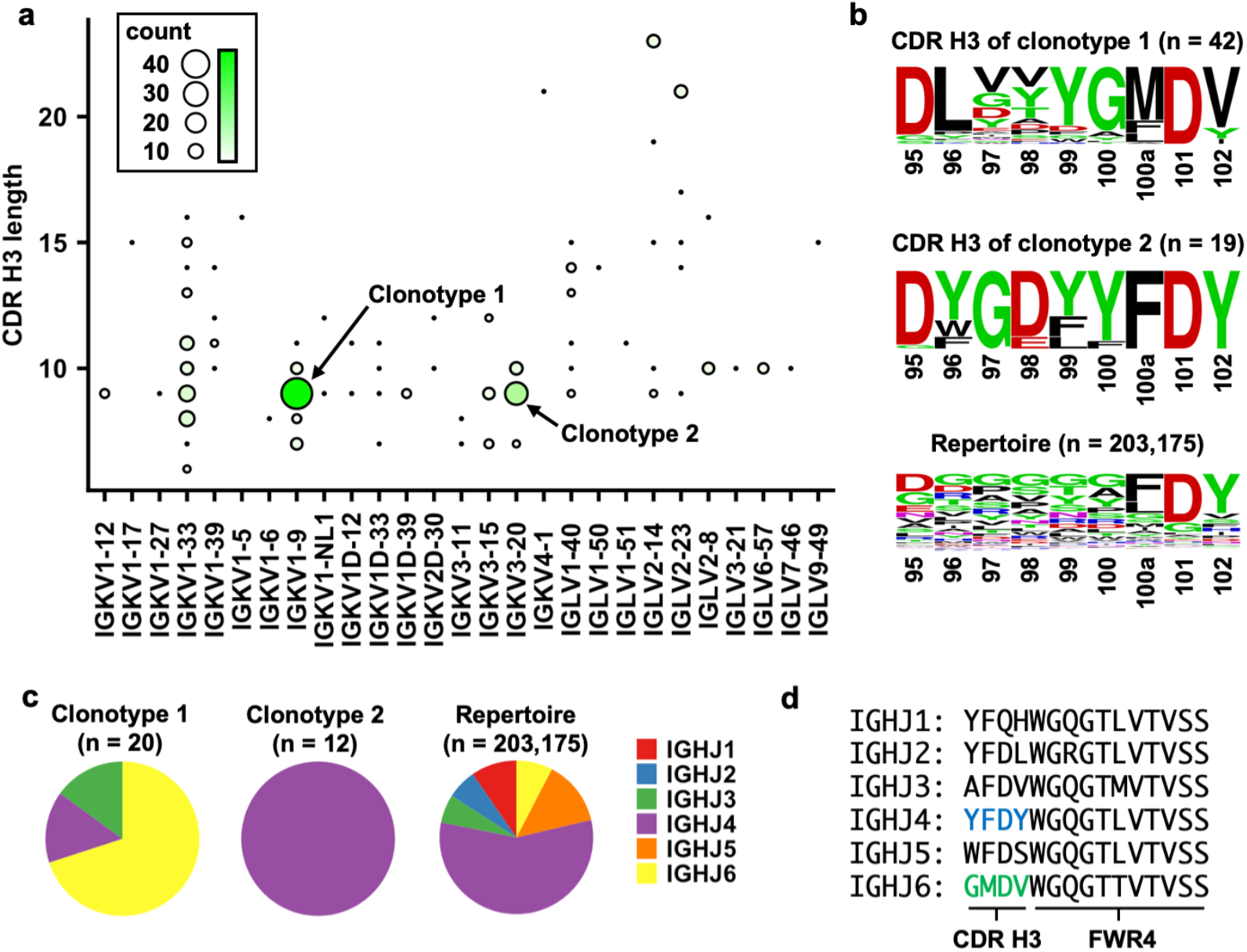
Two major clonotypes of IGHV3-53/3-66 antibodies to SARS-CoV-2 RBD. **(a)** The number of IGHV3-53/3-66 RBD antibodies that use the same light chain with the same CDR H3 are tabulated. The two most common combinations are IGKV1-9 pairing with 9 aa CDR H3 and IGKV3-20 pairing with 9 aa CDR H3, denoted as clonotype 1 and clonotype 2, respectively. **(b)** Sequence logos for the CDR H3 regions of IGHV3-53/66 antibodies that pair with IGKV1-9 or IGKV3-20. A sequence logo for the CDR H3 regions of 203,175 IGHV3-53/3-66 antibodies from Observed Antibody Space database^83^ that have a CDR H3 length of 9 aa is shown for reference (repertoire). The position of each residue is labeled on the x-axis based on Kabat numbering. **(c)** IGHJ gene usage for clonotypes 1 and 2 as well as 203,175 IGHV3-53/3-66 antibodies from Observed Antibody Space database that have a CDR H3 length of 9 aa (repertoire) are shown as pie charts. For antibodies in clonotypes 1 and 2, only those with nucleotide sequence information available were analyzed. **(d)** Amino acid sequences for different IGHJs are shown.

We further determined IGHJ gene usage in the two major clonotypes of IGHV3-53/3-66 RBD antibodies. Among the IGHV3-53/3-66 RBD antibodies with a CDR H3 length of 9 amino acids, we observed a statistically significant bias in IGHJ gene usage (p-value = 2e-6, Fisher’s exact test), where clonotypes 1 and 2 preferentially pair with IGHJ6 and IGHJ4, respectively (Figure 1c). In fact, IGHJ6 encodes the last four amino acids (GMDV) in CDR H3 that are highly conserved in clonotype 1 (Figure 1d, Supplementary Figure 1a). Similarly, IGHJ4 encodes the last four amino acids (YFDY) in CDR H3 that are highly conserved in clonotype 2 (Figure 1d, Supplementary Figure 1b). Taken together, we demonstrate that IGHV3-53/3-66 RBD antibodies can be categorized into at least two public clonotypes.

### Structural analysis of signature motifs on CDR H3

We further investigated sequence signatures of CDR H3s in clonotypes 1 and 2 (Figure 1b). In particular, we focused on amino acid residues 96, 98 and 100 in CDR H3 since these residues show clear patterns of differential amino-acid preference between clonotype 1 and clonotype 2 antibodies. Subsequently, analysis was performed on structures of BD-604 (PDB 7CH4) and CC12.1 (PDB 6XC2), which are two clonotype 1 antibodies, as well as BD-629 (PDB 7CH5) and CC12.3 (PDB 6XC4), which are two clonotype 2 antibodies.

Residue 96 is usually Leu in clonotype 1 antibodies, while an aromatic residue, usually Tyr, occupies residue 96 in clonotype 2 antibodies. While V_H_ L96 interacts with Y489 of the RBD in clonotype 1 antibodies via van der Waals interactions, V_H_ F/Y96 is located at the center of a π-π stacking network that involves F456, Y489 and V_H_ Y100 (Figure 2a, 2b, Supplementary Figure 2a, 2b; left panels). Substituting V_H_ L96 in clonotype 1 with Y96 would result in a clash with RBD Y489, whereas substituting V_H_ F/Y96 in clonotype 2 with L96 would abolish the π-π stacking network but still maintain a hydrophobic core.

**Figure 2.**
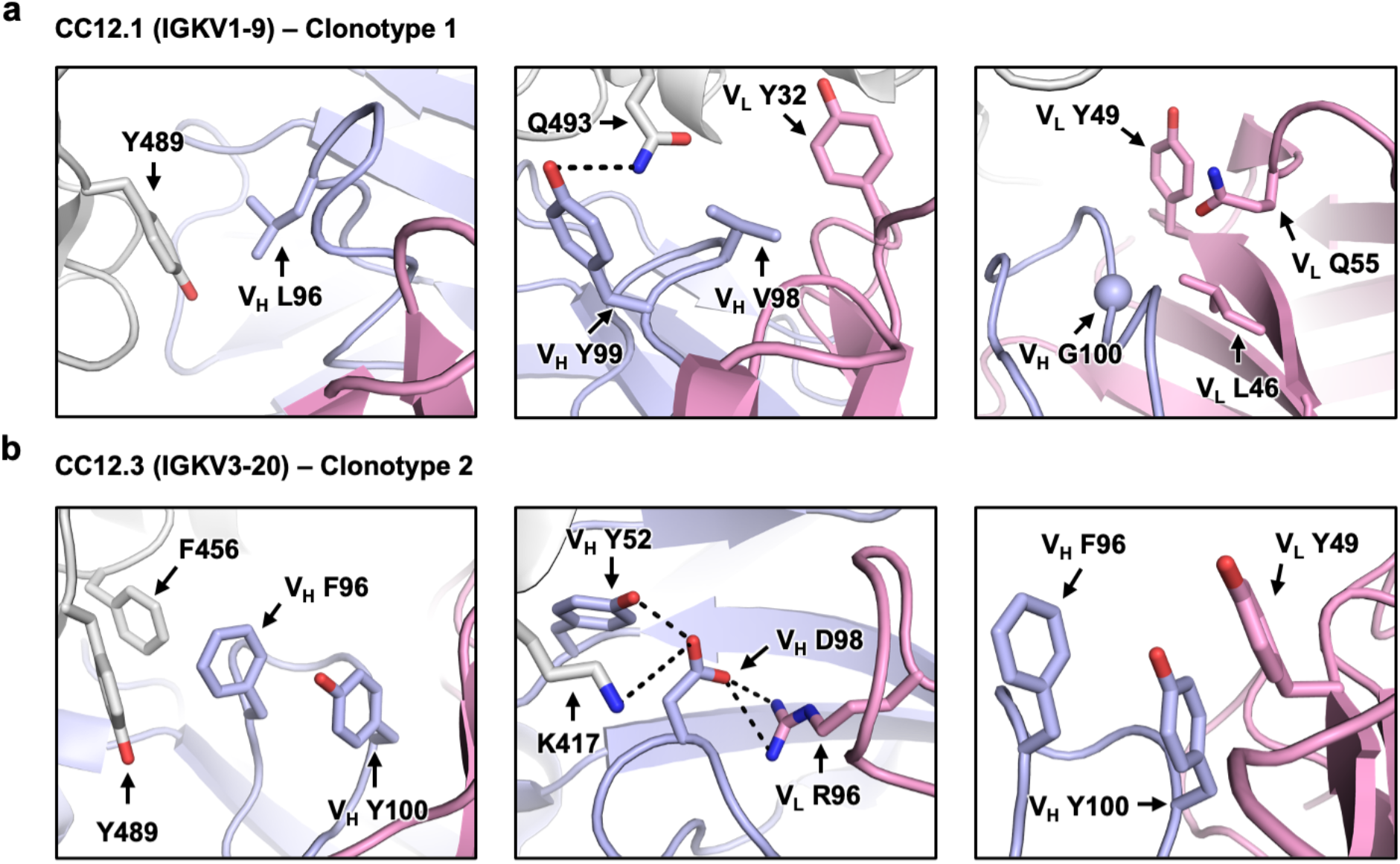
Structural analysis of sequence signatures in CDR H3 of clonotypes 1 and 2. **(a)** Interaction of L96, V98 and G100 (Kabat numbering) in CDR H3 of CC12.1 (PDB 6XC2) with the IGKV1-9 light chain of the antibody, and SARS-CoV-2 RBD. **(b)** Interaction of F96, D98 and Y100 (Kabat numbering) in CDR H3 of CC12.3 (PDB 6XC4) with the IGKV3-20 light chain of the antibody, and SARS-CoV-2 RBD. Gray: RBD; Light blue: heavy chain; Pink: light chain.

Residue 98 in CDR H3 of clonotype 1 antibodies does not show a strong amino-acid preference, since it is located in a relatively open space (Figure 1b, 2a, Supplementary Figure 2a; middle panels). On the other hand, a highly conserved acidic residue at position 98 in the CDR H3 loop of clonotype 2 antibodies contributes to formation of hydrogen bond interactions with V_H_ Y52 as well as electrostatic interactions with RBD K417 and V_L_ R96 (Figure 2b, Supplementary Figure 2b; middle panels). Consistently, V_L_ R96 is highly conserved in clonotype 2 antibodies, but not in other IGHV3-53/3-66 RBD antibodies (Supplementary Figure 3). Thus, the electrostatic interactions between V_H_ D/E98 and V_L_ R98 are highly conserved in clonotype 2 antibodies and can likely help stabilize the CDR H3 loop conformation to minimize entropic cost upon binding to SARS-CoV-2 RBD.

Residue 100 is usually Gly in CDR H3 of clonotype 1 antibodies (Figure 1b). Structural analysis shows that small, non-polar amino acids are favored at position 100 due to the limited space around that residue (Figure 2a, Supplementary Figure 2a; right panels). Moreover, G100 in clonotype 1 has a positive Φ angle, which is typically less favorable for non-Gly amino acids. In contrast, residue 100 is a highly conserved Tyr in CDR H3 of clonotype 2 antibodies (Figure 1b). Structural analysis shows that V_H_ Y100 contributes to the π-π stacking network that is formed via the aromatic ring at V_H_ residue 96 (see above) and an aromatic residue at V_L_ residue 49 (Figure 2b, Supplementary Figure 2b; right panels).

Additionally, we investigated the structural basis of the conservation of V_H_ Y102 among clonotype 2 antibodies. Structural analysis reveals that V_H_ Y102 interacts with RBD Y486 via π-π interactions (Supplementary Figure 4). Only IGHJ4 offers a bulky aromatic side chain at residue 102 (Figure 1d), which explains the common usage of IGHJ4 in clonotype 2 antibodies. In contrast, clonotype 1 antibodies frequently use IGHJ6 (Figure 1d), which has a much shorter Val at residue 102, most likely because IGHJ6 encodes a Gly at residue 100 that can avoid steric clashes with the light chain (see above, Figure 2a, Supplementary Figure 2a; right panels). Of note, the only other IGHJ gene that encodes a non-bulky amino acid at residue 100 is IGHJ3 (Ala). IGHJ1, IGHJ2, IGHJ4, and IGHJ5 all encode a bulky residue at residue 100 (Figure 1d), which may be disfavored in clonotype 1 antibodies due to the limited space where V_H_ residue 100 is located (Supplementary Figure 5). Overall, our structural analyses provide a structural basis for the differential signature sequence motifs in CDR H3 between clonotype 1 and clonotype 2 antibodies.

### Incompatibility of CDR H3 between clonotype 1 and clonotype 2 antibodies

To understand the influence of light-chain usage in CDR H3 sequences, we performed a structural alignment of RBD-bound CDR H3 from two clonotype 1 antibodies, namely BD-604 and CC12.1, and two clonotype 2 antibodies, namely BD-629 and CC12.3 (Supplementary Figures 2c-2f). While the CDR H3 conformations are similar within each clonotype (RMSD ranges from 0.27 to 0.41 Å), they are quite different between clonotypes (RMSD ranges from 0.77 Å to 1.5 Å). Although our sample size is small, this analysis suggests that antibodies from clonotypes 1 and 2 have different preferences for their CDR H3 conformations. Such differential preference of CDR H3 conformations may be partly influenced by light-chain usage, as indicated by the structural analyses above on V_H_ residues 96, 98, and 100 (Figure 2, Supplementary Figures 2 and 5).

To experimentally examine the compatibility between CDR H3 and the light chains from clonotype 1 and clonotype 2 antibodies, we focused on antibodies COV107-23 (clonotype 1) and COVD21-C8 (clonotype 2). The heavy-chain sequences of these two antibodies only differ by four amino acids in CDR H3, namely V_H_ residues 96, 98, 99, and 100 (Supplementary Figure 6a). Of note, COV107-23 uses IGHJ4, which is seldom observed among clonotype 1 antibodies but highly preferred in clonotype 2 antibodies (Figure 1c), to encode the two amino acids at the C-terminus of its CDR H3 (Supplementary Figure 6b). Both COV107-23 and COVD21-C8 bind strongly to the SARS-CoV-2 RBD, with dissociation constants (K_D_) of 1 nM and 4 nM, respectively (Figure 3a). However, when their light chains are swapped, their binding affinity to the RBD is weakened substantially to K_D_ > 1 μM. We further determined apo crystal structures of COV107-23 paired with its native light chain and with the light chain from COVD21-C8 to 2.0 Å and 3.3 Å, respectively (Supplementary Table 2). The conformations of CDR H3 indeed differ when paired with different light chains, as exemplified by the 3.3 Å displacement of V_H_ G97 near the tip of CDR H3 and different side-chain orientations of V_H_ T98 (Figure 3b). In addition, a type I’ β-turn is observed at the tip of CDR H3 in COV107-23 when paired with its native light chain but not with the light chain from COVD21-C8 (Figure 3c). These observations demonstrate that the conformation of CDR H3 changes substantially when IGKV1-9 in COV10-23 is swapped to IGKV3-20, which abolishes the binding to RBD (Figure 3a). The CDR H3 conformation is therefore a determinant for compatibility between the CDR H3 sequence and the light chain in IGHV3-53/3-66 RBD antibodies.

**Figure 3.**
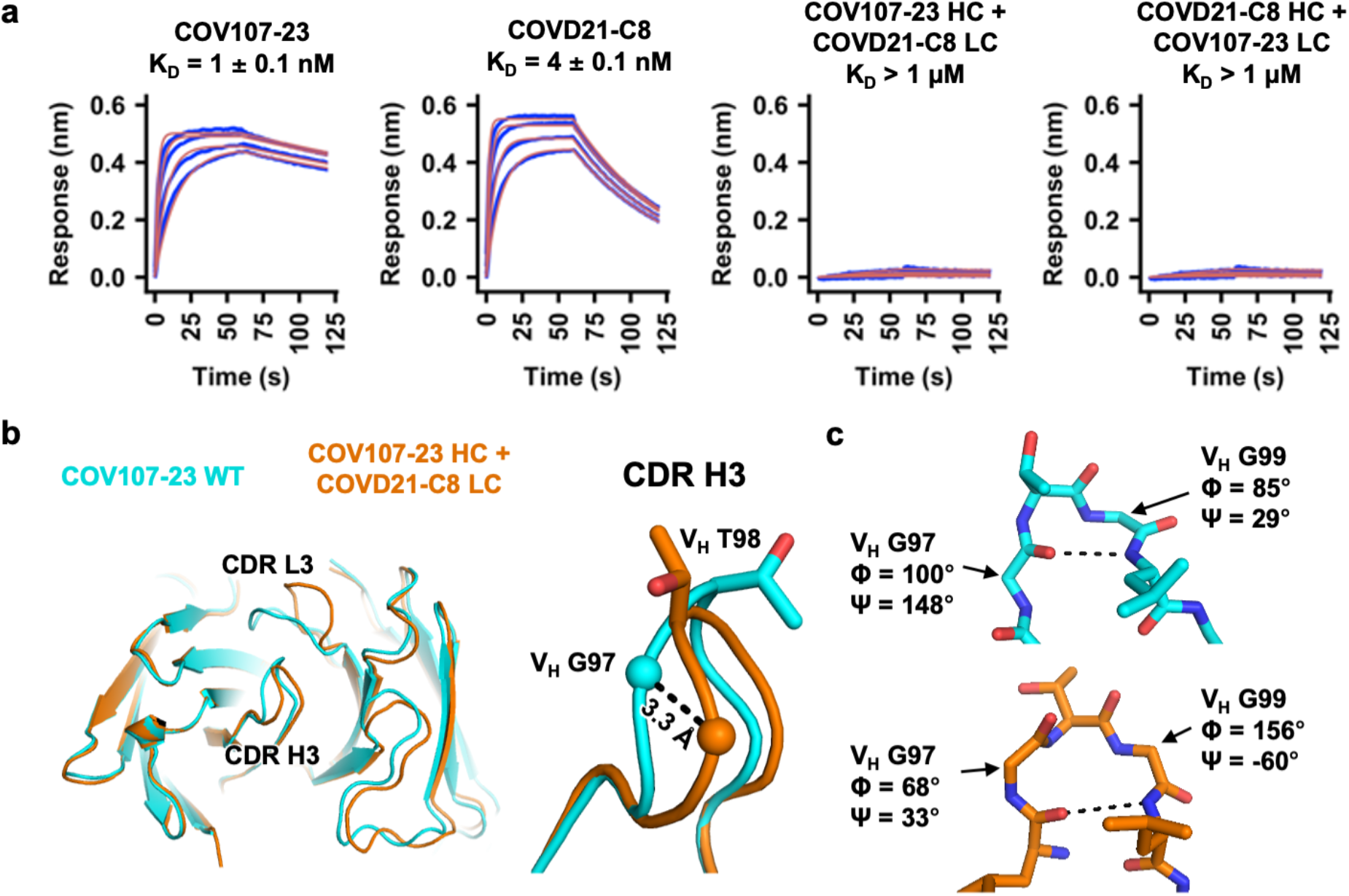
Specific pairing of CDR H3 and light chain is critical for IGHV3-53/3-66 antibody binding to SARS-CoV-2 RBD. **(a)** Binding of different Fabs to SARS-CoV-2 RBD was measured by biolayer interferometry with RBD loaded onto the biosensor and Fab in solution. Y-axis represents the response. Dissociation constant (K_D_) for each Fab was obtained using a 1:1 binding model, which is represented by the red curves. **(b)** Fab crystal structures of wild-type (WT) COV107-23 and COV107-23 heavy chain pairing with COVD21-C8 light chain are compared. Left panel: structural alignment using residues 1-90 of the heavy chain. Right panel: Zoom-in view for the CDR H3. **(c)** Conformations at the tips of the CDR H3s in WT COV107-23 and COV107-23 heavy chain pairing with COVD21-C8 light chain are shown. A β-turn is observed in the CDR H3 of WT COV107-23, with V_H_ G97 and V_H_ G99 at i and i+2 positions, respectively.

### Compatibility of different CDR H3 variants with IGHV1-9 for binding to RBD

Besides antibodies from clonotypes 1 and 2, other IGHV3-53/3-66 RBD antibodies with a range of CDR H3 lengths pair with different light chains (Figure 1a). We further aimed to expand our analysis on CDR H3 compatibility to include CDR H3 from IGHV3-53/3-66 RBD antibodies other than clonotypes 1 and 2. In particular, we focused on identifying CDR H3 sequences that are compatible with IGKV1-9, which is used by clonotype 1 antibodies for binding to RBD. We first compiled a list of 143 CDR H3 variants that were observed in IGHV3-53/3-66 RBD antibodies (Supplementary Table 3). A yeast display library was then constructed with these 143 CDR H3 variants in the B38 antibody, which is a IGHV3-53/IGKV1-9 RBD antibody^26^. Subsequently, fluorescence-activated cell sorting (FACS) was performed on the yeast display library based on antibody expression level and binding to SARS-CoV-2 RBD (Supplementary Figures 7 and 8). The enrichment level of each CDR H3 variant in the sorted library was quantified by next-generation sequencing (see Methods, Supplementary Table 4). CDR H3 variants that were positively enriched in binding (log_10_ enrichment > 0) are derived from both IGKV1-9 and non-IGKV1-9 antibodies (Figure 4a). The native CDR H3 for B38 has a log_10_ enrichment level of - 0.002. As a result, positively enriched CDR H3 variants should have a higher affinity than wild-type B38. A total of 68% (17 out of 25) binding-enriched CDR H3 variants have a length of 9 amino acids, whereas only 31% (37 out of 118) have a length of 9 amino acids in the non-enriched group (Figure 4b). Interestingly, binding-enriched CDR H3 variants with a length of 9 amino acids displayed very similar sequence features as that of clonotype 1 antibodies obtained from literature mining (Figure 1b and 4c). Of note, 41% (7 out of 17) binding-enriched CDR H3 variants with a length of 9 amino acids come from non-IGKV1-9 antibodies. Overall, our yeast display screen indicates that certain CDR H3s from non-IGKV1-9 RBD antibodies are compatible with IGKV1-9 for RBD binding and have similar sequence features as those CDR H3s from clonotype 1 antibodies.

**Figure 4.**
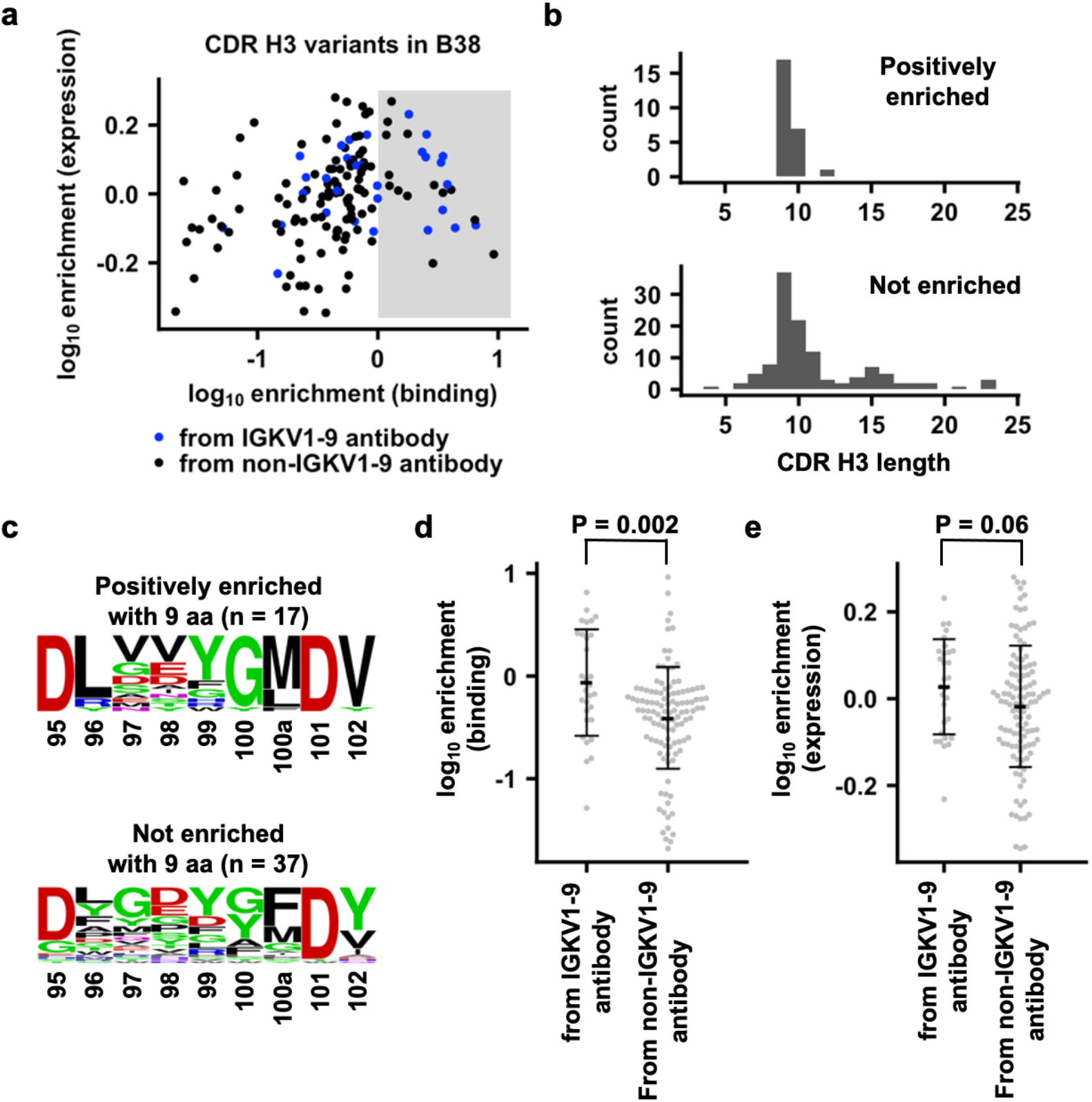
Binding and expression profiling of 143 CDR H3 variants in B38 antibody. **(a)** For each of the 143 CDR H3 variants, the enrichment in occurrence frequencies after FACS selections for binding to RBD and expression level are shown. Blue: CDR H3 variants that are derived from IGHV3-53/3-66 RBD antibodies that use IGKV1-9. Black: CDR H3 variants that are derived from IGHV3-53/3-66 RBD antibodies that do not use IGKV1-9. Shaded area indicates log_10_ enrichment in binding > 0. **(b)** The amino-acid length distribution of CDR H3 variants that are positively enriched in binding (log_10_ enrichment in binding > 0) or not (log_10_ enrichment in binding ≤ 0) is shown. **(c)** Sequence logos are shown for CDR H3 variants with 9 aa (Kabat numbering) that are positively enriched or not enriched. **(d)** Comparison of log_10_ enrichment in binding for CDR H3 variants from IGHV3-53/3-66 RBD antibodies that use IGKV1-9 and those that do not use IGKV1-9. **(e)** Comparison of log_10_ enrichment in expression for CDR H3 variants from IGHV3-53/3-66 RBD antibodies that use IGKV1-9 and those that do not use IGKV1-9. **(d-e)** Student’s t-test was used to compute the p-value.

We noticed that some CDR H3 sequences that come from IGKV1-9 RBD antibodies do not enrich in binding. One possibility is that they are still able to bind to RBD, but with a lower affinity than B38, which has a K_D_ of 70 nM to the RBD^26^. However, as shown by our yeast display screen, CDR H3 sequences from IGKV1-9 antibodies in general have a significantly stronger binding to RBD than those from non-IGKV1-9 antibodies (p-value = 0.002, Figure 4d), whereas their expression level is only marginally higher than that from non-IGKV1-9 antibodies (p-value = 0.06, Figure 4d).

### Y58F is a signature SHM in IGHV3-53/3-66 RBD antibodies

We further aimed to understand if there are common SHMs among IGHV3-53/3-66 RBD antibodies. We first categorized IGHV3-53/3-66 RBD antibodies from convalescent SARS-CoV-2 patients by CDR H3 length. The occurrence frequencies of individual SHMs in each category were then analyzed (Figure 5a). This analysis included 214 IGHV3-53/3-66 RBD antibodies that have sequence information available. One clear observation is that Y58F is highly common among IGHV3-53/3-66 RBD antibodies with a CDR H3 length of less than 15 amino acids, but completely absent when the CDR H3 length is 15 amino acids or above, suggesting that Y58F improves the binding of affinity IGHV3-53/3-66 antibodies to RBD only when they have a short CDR H3 loop (CDR H3 < 15 amino acids). To understand the effect of Y58F on the binding affinity of IGHV3-53/3-66 antibodies to the RBD, we compared the binding affinity of the same antibodies that carry either Y58 or F58 to the RBD. In particular, we focused on three IGHV3-53/3-66 RBD antibodies that have a CDR H3 length of 9 amino acids – one in clonotype 1 (COV107-23), and two in clonotype 2 (COVD21-C8 and CC12.3). Our BLI experiments showed that the Y58F mutation dramatically improved the affinity of the three antibodies (COV107-23, COVD21-C8 and CC12.3) by ~10-fold to ~1000-fold (Figure 5b, Supplementary Figure 9). As a control, we also performed the same experiment on an IGHV3-53/3-66 antibody with a CDR H3 length of 15 amino acids, namely COVA2-20. In contrast to those three IGHV3-53/3-66 RBD antibodies with a short CDR H3, COVA2-20 shows similar binding affinity to RBD between Y58 and F58 variants (Figure 5b, Supplementary Figure 5). Taken together, our results show that Y58F appears to be a signature SHM in IGHV3-53/3-66 RBD antibodies with CDR H3 length of < 15 amino acids. In fact, the results here are consistent with our previous finding that IGHV3-53/3-66 RBD antibodies with CDR H3 length of 15 amino acids or longer adopt a different binding mode as compared to those with a shorter CDR H3^54^.

**Figure 5.**
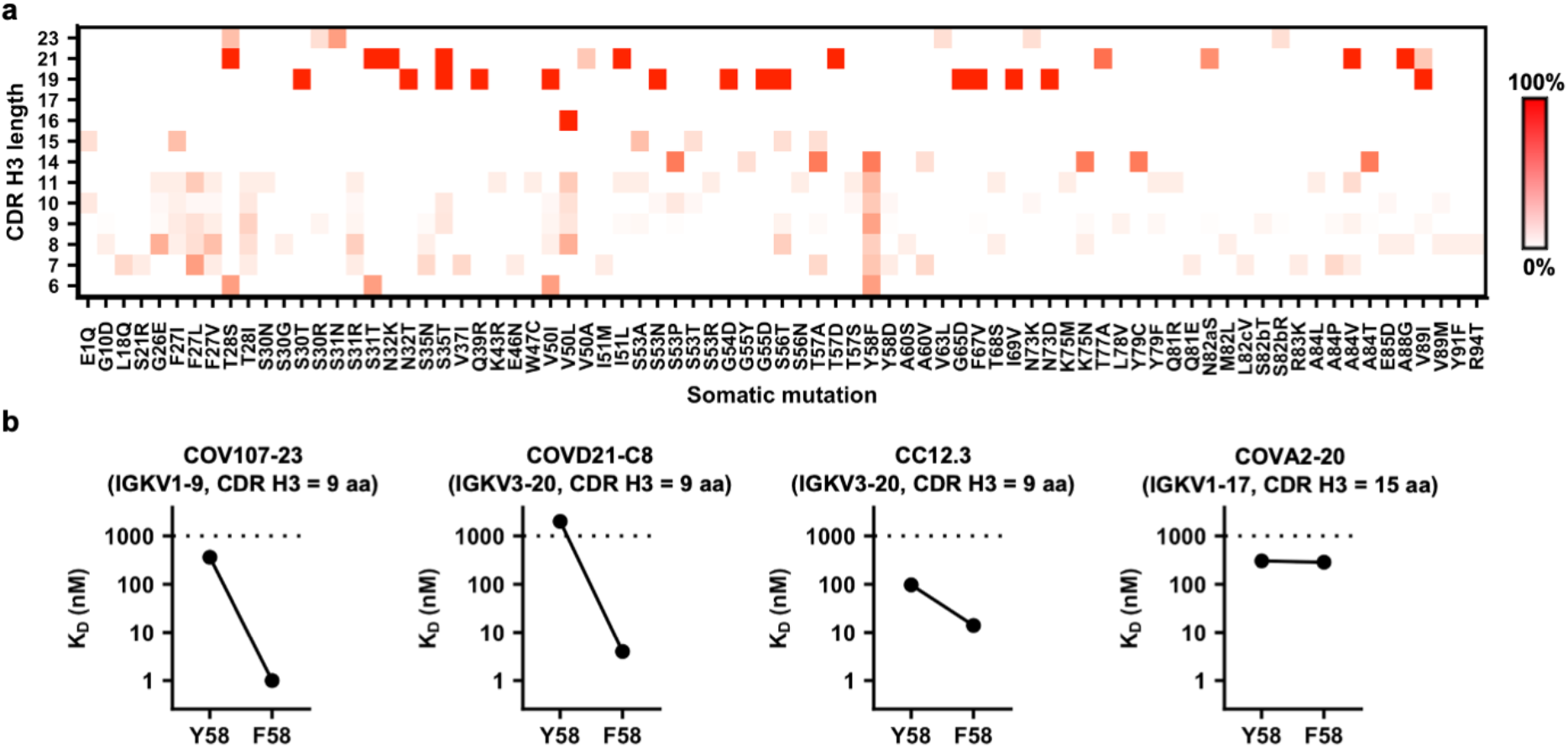
Y58F is a signature somatic hypermutation in IGHV3-53/3-66 RBD antibodies with a short CDR H3. **(a)** IGHV3-53/3-66 RBD antibodies are categorized based on their CDR H3 length (Kabat numbering). Occurrence frequencies of individual somatic hypermutations in different categories were quantified and shown as a heatmap. **(b)** Both Y58 and F58 variants were constructed for four IGHV3-53 antibodies. Binding affinity (K_D_) of each of these antibodies as Fab format to SARS-CoV-2 RBD was measured by biolayer interferometry with RBD loaded on the biosensor and Fab in solution. Y-axis represents the response. Dissociation constants (K_D_) for the Fabs were obtained using a 1:1 binding model. Of note, the WTs of COV107-23, COVD21-C8, and CC12.3 contain F58, whereas the WT of COVA2-20 contains Y58.

Interestingly, a Y58F mutation results in a loss of hydrogen bonding interactions between residue 58 of the heavy chain and T415 of the RBD (Supplementary Figure 10), yet the mutation significantly increases the binding affinity of the antibody to the RBD. We then performed a structural analysis on seven IGHV3-53/66 RBD antibodies with Y58F mutation and nine without^26,29,38,40,47,54-57^. Our results indicate that, by removal of the hydroxyl group, the side chain of Y58F moves closer to the backbone carbon of RBD T415 (Supplementary Figure 10). The average distance between the centroid of the side-chain aromatic ring at V_H_ residue 58 and the backbone carbon of RBD T415 are 5.3 Å and 5.9 Å for antibodies that carry F58 and Y58, respectively. Since T-shaped π-π stacking is optimal at around 5.0 to 5.2 Å^58,59^, F58 but not Y58 can form strong T-shaped π-π stacking interactions with the amide backbone of RBD T415. This observation can at least partly explain why Y58F improves affinity despite the loss of a hydrogen bond with the RBD.

## Discussion

While several studies to date have described IGHV3-53/3-66 as a commonly used germline for SARS-CoV-2 RBD antibodies^23,31,47,54^, the exact sequence requirements for generating an IGHV3-53/3-66 antibody to SARS-CoV-2 RBD has remained largely elusive. As a result of numerous efforts from multiple groups in isolating RBD antibodies and reporting their sequences^20–40^, detailed characterization of RBD antibody sequence features has become possible. Through sequence analysis, biophysical experiments, and high-throughput screening, we identified distinct sequence requirements for two public clonotypes (clonotypes 1 and 2) of IGHV3-53/3-66 RBD antibodies. In fact, the frequent occurrence of IGHV3-53/3-66 RBD antibodies with IGHJ6 and a CDR H3 length of 9 amino acids, which are germline features of clonotype 1 antibodies, have also been reported in previous publications^23,60^.

One important finding in this study is that the CDR H3 sequence that supports IGHV3-53/3-66 antibodies binding to RBD is light chain-dependent. This finding is consistent with our previous observation that there is a large diversity of CDR H3 sequences in IGHV3-53/3-66 RBD antibodies^54^. In addition, our findings explain a recent observation by Banach and colleagues^61^ who showed that swapping the heavy and light chains of different IGHV3-53/3-66 RBD antibodies often substantially reduced their neutralization potency. Therefore, IGHV3-53/3-66 provides a robust framework to generate different public clonotypes that have distinct CDR H3 and light chain sequence signatures. While only two major clonotypes of IGHV3-53/3-66 RBD antibodies are examined in this study, it will be worth characterizing other minor clonotypes to obtain a more complete understanding of the compatibility between CDR H3 sequence and light-chain identity among IGHV3-53/3-66 RBD antibodies.

Although this study revealed that Y58F is a common SHM that improves the affinity of IGHV3-53/3-66 antibodies with a short CDR H3 to RBD, other common SHMs have also shown up in our sequence analysis (Figure 5a), albeit with a lower frequency. Most noticeably, a cluster of common SHMs is found in V_H_ framework region 1 from residues 26 to 28. This cluster of SHMs is also likely to be important for affinity maturation to RBD. A recent study has indeed shown that SHMs V_H_ F27V and T28I together increase affinity by 100-fold of an IGHV3-53/3-66 antibody to the SARS-CoV-2 RBD^38^. Additional common SHMs among IGHV3-53/3-66 RBD antibodies with a short CDR H3 include S31R in CDR H1 and V50L in CDR H2 (Figure 5a). As a result, while IGHV3-53/3-66 RBD antibodies do not require any SHM to neutralize SARS-CoV-2 ^57^, this study along with others have shown that SHM can substantially improve the binding affinity of IGHV3-53/3-66 antibodies to RBD^38,57^. Consistently, RBD antibodies from convalescent SARS-CoV-2 patients have significantly more SHMs and higher neutralization potency at 6 months post-infection than at 1-month post-infection^62^.

Circulating SARS-CoV-2 mutant variants represent a major ongoing challenge to natural immunity and vaccination. In particular, a lot of attention has been focused on RBD mutation E484K, which has emerged in multiple independently SARS-CoV-2 lineages^63,64^ and can alter the antigenicity of the spike protein^65–67^. Another naturally occurring RBD mutation, K417N, which has emerged in South Africa and Brazil (B.1.351 lineage and B.1.1.28, respectively)^63,64,68^, has recently been shown to also alter antigenicity of the spike protein^66,69-71^. Consistently, we found that K417N dramatically decreased the binding of COV107-23 (clonotype 1) and COVD21-C8 (clonotype 2) to RBD (Supplementary Figures 11a-11b). In fact, K417 forms an electrostatic interaction with the signature residue V_H_ D/E98 of CDR H3 in clonotype 2 antibodies (Figure 2b) and can also interact with CDR H3 of clonotype 1 antibodies (Supplementary Figure 11c), providing a structural explanation for its change in antigenicity. Constant antigenic drift of SARS-CoV-2 is unavoidable if it keeps circulating among humans. Thus, sustained efforts in characterizing the antibody response to SARS-CoV-2 as it evolves will not only benefit vaccine development and assessment, but also improve our fundamental understanding of the ability of the antibody repertoire to rapidly respond to viral infections.

## Methods

### Literature mining for antibodies to SARS-CoV-2 RBD

Sequences of anti-SARS-CoV-2 RBD from convalescent patients infected with SARS-CoV-2 were obtained from published articles^20–40^ (Supplementary Table 1). IgBlast was used to identify somatic hypermutations and analyze IGHJ gene usage^72^. Of note, IgBlast can only identify IGHJ gene usage for antibodies with available nucleotide sequences. Sequence logos were generated by WebLogo^73^.

### Expression and purification of Fc-tagged RBD

The receptor-binding domain (RBD) (residues 319-541) of the SARS-CoV-2 spike (S) protein (GenBank: QHD43416.1) was fused with an N-terminal Igk secretion signal and a C-terminal SSSSG linker followed by an Fc tag and cloned into a phCMV3 vector. The plasmid was transiently transfected into Expi293F cells using ExpiFectamine™ 293 Reagent (Thermo Fisher Scientific) according to the manufacturer’s instructions. The supernatant was collected at 7 days post-transfection. The Fc-tagged RBD was purified with by KanCapA protein A affinity resin (Kaneka).

### Expression and purification of Fabs

Fab heavy and light chains were cloned into phCMV3. Heavy chain Y58F or F58Y mutants were constructed using the QuikChange XL Mutagenesis kit (Stratagene) according to the manufacturer’s instructions. The plasmids were transiently co-transfected into Expi293F cells at a ratio of 2:1 (HC:LC) using ExpiFectamine™ 293 Reagent (Thermo Fisher Scientific) according to the manufacturer’s instructions. The supernatant was collected at 7 days post-transfection. The Fab was purified with a CaptureSelect™ CH1-XL Pre-packed Column (Thermo Fisher Scientific).

### Biolayer interferometry binding assay

Binding assays were performed by biolayer interferometry (BLI) using an Octet Red96e instrument (FortéBio) as described previously^74^. Briefly, Fc-tagged SARS-CoV-2 RBD proteins at 20 to 100 μg/ml in 1x kinetics buffer (1x PBS, pH 7.4, 0.01% w/v BSA and 0.002% v/v Tween 20) were loaded onto streptavidin (SA) biosensors and incubated with the indicated concentrations of Fabs. The assay consisted of five steps: 1) baseline: 60 s with 1x kinetics buffer; 2) loading: 300 s with His6-tagged S or RBD proteins; 3) baseline: 60 s with 1x kinetics buffer; 4) association: 60 s with samples (Fab or IgG); and 5) dissociation: 60s with 1x kinetics buffer. For estimating the exact K_D_, a 1:1 binding model was used.

### X-ray crystallography

Fabs COV107-23 (15 mg/ml) and COV107-23 paired with the light chain of COVD21-C8 (COV107-23-swap, 14 mg/ml) were screened for crystallization using the 384 conditions of the JCSG Core Suite (Qiagen) on our custom-designed robotic CrystalMation system (Rigaku) at Scripps Research by the vapor diffusion method in sitting drops containing 0.1 μl of protein and 0.1 μl of reservoir solution. For COV107-23, optimized crystals were grown in 0.085 M of sodium citrate - citric acid pH 5.6, 0.17 M ammonium acetate, 15% (v/v) glycerol, and 25.5% (w/v) polyethylene glycol 4000 at 20°C. For COV107-23-swap, optimized crystals were grown in 0.1 M of sodium citrate pH 4, 1 M lithium chloride, and 20% (w/v) polyethylene glycol 6000 at 20°C. Crystals were grown for 7 days and then harvested and flash cooled in liquid nitrogen. Diffraction data were collected at cryogenic temperature (100 K) at Stanford Synchrotron Radiation Lightsource (SSRL) on the Scripps/Stanford beamline 12-1 with a beam wavelength of 0.97946 Å, and processed with HKL2000^75^. Structures were solved by molecular replacement using PHASER^76^, where the models were generated by Repertoire Builder (https://sysimm.org/rep_builder/)^77^. Iterative model building and refinement were carried out in COOT^78^ and PHENIX^79^, respectively.

### Construction of plasmids and CDR H3 library

143 oligonucleotides (Supplementary Table 3) encoding CDR H3 were obtained from Integrated DNA Technologies (IDT) and PCR-amplified using 5’-ACC TAC AGA TGA ATT CTC TTA GGG CAG AAG ATA CCG CCG TCT ACT ACT GC-3’ as forward primer and 5’-GGG CCT TTT GTA GAA GCT GAA CTC ACA GTG ACG GTA GTC CCT TGT CCC CA-3 as reverse primer. Then, the amplified oligonucleotide pool was gel-purified using a GeneJET Gel Extraction Kit (Thermo Scientific).

Wild-type (WT) B38 yeast display plasmid, pCTcon2_B38, was generated by cloning the coding sequence of (from N-terminal to C-terminal, all in-frame) Aga2 secretion signal, B38 Fab light chain, V5 tag, ERBV-1 2A self-cleaving peptide, Aga2 secretion signal, B38 Fab heavy chain, HA tag, and Aga2p, into the pCTcon2 vector^80^. pCTcon2_B38 was PCR-amplified using 5’-TGG GGA CAA GGG ACT ACC GTC ACT GTG-3 as forward primer and 5-GCA GTA GTA GAC GGC GGT ATC TTC TGC-3’ as reverse primer to generate the linearized vector. The PCR product was then gel-purified.

### Yeast antibody display library generation

5 μg of the amplified oligonucleotide pool and 4 μg of purified linearized vector were transformed into *Saccharomyces cerevisiae* EBY100 via electroporation following previously published protocol^81^ to generate a B38 yeast display library with different CDR H3 variants.

### Fluorescence-activated cell sorting of yeast antibody display library

100 μl of WT B38 yeast antibody display library glycerol stock was recovered in 50 ml SD-CAA medium (2% w/v D-glucose, 0.67% w/v yeast nitrogen base with ammonium sulfate, 0.5% w/v casamino acids, 0.54% w/v Na_2_HPO_4_, 0.86% w/v NaH_2_PO_4_·H_2_O, all dissolved in deionized water) by incubating at 27°C with shaking at 250 rpm until OD_600_ reached between 1.5 and 2.0. At this time, 15 ml of the yeast culture was harvested, and the yeast pellet was obtained via centrifugation at 4,000 × *g* at 4°C for 5 min. The supernatant was discarded, and SGR-CAA (2% w/v galactose, 2% w/v raffinose, 0.1% w/v D-glucose, 0.67% w/v yeast nitrogen base with ammonium sulfate, 0.5% w/v casamino acids, 0.54% w/v Na_2_HPO_4_, 0.86% w/v NaH_2_PO_4_·H_2_O, all dissolved in deionized water) was added to make up the volume to 50 ml. The yeast culture was then transferred to a baffled flask and incubated at 18°C with shaking at 250 rpm. Once OD_600_ has reached between 1.3 and 1.6, 1 ml of yeast culture was harvested, and the yeast pellet was obtained via centrifugation at 4,000 × *g* at 4°C for 5 min. The pellet was subsequently washed with 1 ml of 1x PBS twice. After the final wash, cells were resuspended in 1 ml of 1x PBS.

Then, for expression assay, 1 μg of PE anti-HA.11 (epitope 16B12, BioLegend, Cat. No. 901517) buffer-exchanged into 1x PBS was added to the cells. A negative control was set up with nothing added to the PBS-resuspended cells. Samples were incubated overnight at 4°C with rotation. Then, the yeast pellet was washed twice in 1x PBS and resuspended in FACS tubes containing 2 ml 1X PBS. Using a BD FACS Aria II cell sorter (BD Biosciences), PE-positive cells were collected in 1 ml of SD-CAA containing 1x Penicillin/Streptomycin. Cells were then collected via centrifugation at 4,500 rpm at 20°C for 15 min. The supernatant was discarded. Subsequently, the pellet was resuspended in 100 μl of SD-CAA and plated on SD-CAA plates at 37°C. After 40 h, colonies were collected in 2 ml of SD-CAA. Frozen stocks were made by reconstituting the pellet in 15% v/v glycerol (in SD-CAA medium) and then stored at −80°C.

For binding assay, 20 μg of SARS-CoV-2 S RBD-Fc was added to washed cells. A negative control was set up with nothing added to the PBS-resuspended cells. Samples were incubated overnight at 4°C with rotation. The yeast pellet was then washed twice in 1x PBS. After the last wash, cells were resuspended in 1 ml of 1x PBS. Subsequently, 1 μg of PE anti-human IgG Fc antibody (clone HP6017, BioLegend, Cat. No. 409304) buffer-exchanged into 1x PBS was added to yeast. Cells were incubated at 4°C for 1 h with rotation. The yeast pellet was then washed twice in 1x PBS and resuspended in FACS tubes containing 2 ml 1x PBS. Using a BD FACS Aria II cell sorter (BD Biosciences), PE-positive cells were collected in 1 ml of SD-CAA containing 1x Penicillin/Streptomycin. Cells were then collected via centrifugation at 4,500 rpm at 20°C for 15 min. The supernatant was then discarded. Subsequently, the pellet was resuspended in 100 μl of SD-CAA and plated on SD-CAA plates at 37°C. After 40 h, colonies were collected in 2 ml of SD-CAA, and subsequently pelleted. Frozen stocks were made by reconstituting yeast pellets with 15% v/v glycerol (in SD-CAA medium) and then stored at −80°C.

### Next-generation sequencing of CDR H3 loops

Plasmids from the unsorted yeast display library (input) as well as two replicates of sorted yeast display library based on RBD-binding and expression were extracted from sorted yeast cells using a Zymoprep Yeast Plasmid Miniprep II Kit (Zymo Research) following the manufacturer’s protocol. The CDR H3 region was subsequently amplified via PCR using 5’-ACC TAC AGA TGA ATT CTC TTA GG-3’ and 5’-GGG CCT TTT GTA GAA GCT GAA CT-3’ as forward and reverse primers, respectively. Subsequently, adapters containing sequencing barcodes were appended to the genes encoding the CDR H3 region via PCR. 100 ng of each sample was used for paired-end sequencing using Illumina MiSeq PE150 (Illumina). PEAR was used for merging the forward and reverse reads^82^. Regions corresponding to the CDR H3 were extracted from each paired read. The number of reads corresponding to each CDR H3 variant in each sample is counted. A pseudocount of 1 was added to the final count to avoid division by zero in enrichment calculation. The enrichment for variant *i* was computed as follows:

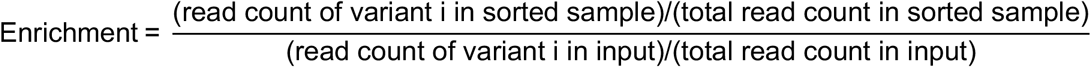

## Code availability

Custom python scripts for analyzing the deep mutational scanning data have been deposited to https://github.com/wchnicholas/IGHV3-53_sequence_features. Files for Rosetta modeling are available at https://github.com/timothyjtan/ighv3-53_3-66_antibody_sequence_features.

## Data availability

Raw sequencing data have been submitted to the NIH Short Read Archive under accession number: BioProject PRJNA691562. The X-ray coordinates and structure factors will be deposited to the RCSB Protein Data Bank prior to publication.

## Acknowledgements

We thank the Roy J. Carver Biotechnology Center at the University of Illinois at Urbana-Champaign for assistance with fluorescence-activated cell sorting and next-generation sequencing. This work was supported by NIH R00 AI139445 (N.C.W.), and Bill and Melinda Gates Foundation INV-004923 (I.A.W.).

## Competing interests

The authors declare no competing interests.

## Author Contributions

T.J.C.T., G.C.P. and N.C.W. conceived and designed the study. J.R.B., M.Y. and B.M.S. expressed and purified the proteins. T.J.C.T., K.K., X.C., J.R.C. and C.B.B. performed the yeast display experiments. T.J.C.T., Y.W. and N.C.W. processed the next-generation sequencing data. M.Y. and X.Z. performed the crystallization, X-ray data collection, determined and refined the X-ray structures. T.J.C.T., M.Y., G.C.P., I.A.W. and N.C.W. analyzed the data. T.J.C.T., M.Y., I.A.W. and N.C.W. wrote the paper and all authors reviewed and/or edited the paper.

## Notes

### Competing Interest Statement

The authors have declared no competing interest.

## References

1 Zhou, P. et al. A pneumonia outbreak associated with a new coronavirus of probable bat origin. Nature 579, 270–273 (2020).

2 Zhu, N. et al. A novel coronavirus from patients with pneumonia in China, 2019. N. Engl. J. Med. 382, 727–733 (2020).

3 Guan, W.-j. et al. Clinical characteristics of coronavirus disease 2019 in China. N. Engl. J. Med. 382, 1708–1720 (2020).

4 Nadim, M. K. et al. COVID-19-associated acute kidney injury: consensus report of the 25th Acute Disease Quality Initiative (ADQI) Workgroup. Nat. Rev. Nephrol. 16, 1–18 (2020).

5 Letko, M., Marzi, A. & Munster, V. Functional assessment of cell entry and receptor usage for SARS-CoV-2 and other lineage B betacoronaviruses. Nat. Microbiol. 5, 562–569 (2020).

6 Wan, Y., Shang, J., Graham, R., Baric, R. S. & Li, F. Receptor recognition by the novel coronavirus from Wuhan: an analysis based on decade-long structural studies of SARS coronavirus. J. Virol. 94, e00127–00120 (2020).

7 Benton, D. J. et al. Receptor binding and priming of the spike protein of SARS-CoV-2 for membrane fusion. Nature 588, 327–330 (2020).

8 Krammer, F. SARS-CoV-2 vaccines in development. Nature 586, 516–527 (2020).

9 Amanat, F. & Krammer, F. SARS-CoV-2 Vaccines: Status Report. Immunity 52, 583–589 (2020).

10 Polack, F. P. et al. Safety and efficacy of the BNT 162b2 mRNA Covid-19 vaccine. N. Engl. J. Med. 383, 2603–2615 (2020).

11 Mahase, E. Covid-19: Moderna vaccine is nearly 95% effective, trial involving high risk and elderly people shows. BMJ: British Medical Journal (Online) 371, doi:https://doi.org/10.1136/bmj.m4471 (2020).

12 Mahase, E. Covid-19: Moderna applies for US and EU approval as vaccine trial reports 94.1% efficacy. BMJ: British Medical Journal (Online) 371, doi:https://doi.org/10.1136/bmj.m4709 (2020).

13 Folegatti, P. M. et al. Safety and immunogenicity of the ChAdOx1 nCoV-19 vaccine against SARS-CoV-2: a preliminary report of a phase 1/2, single-blind, randomised controlled trial. Lancet 396, 467–478 (2020).

14 Ramasamy, M. N. et al. Safety and immunogenicity of ChAdOx1 nCoV-19 vaccine administered in a prime-boost regimen in young and old adults (COV002): a single-blind, randomised, controlled, phase 2/3 trial. Lancet 396, 1979–1993 (2020).

15 Voysey, M. et al. Safety and efficacy of the ChAdOx1 nCoV-19 vaccine (AZD1222) against SARS-CoV-2: an interim analysis of four randomised controlled trials in Brazil, South Africa, and the UK. Lancet 397, 99–111 (2021).

16 Premkumar, L. et al. The receptor binding domain of the viral spike protein is an immunodominant and highly specific target of antibodies in SARS-CoV-2 patients. Sci. Immunol. 5, doi:https://doi.org/10.1126/sciimmunol.abc8413 (2020).

17 Yuan, M., Liu, H., Wu, N. C. & Wilson, I. A. Recognition of the SARS-CoV-2 receptor binding domain by neutralizing antibodies. Biochem. Biophys. Res. Commun., doi:https://doi.org/10.1016/j.bbrc.2020.10.012 (2020).

18 Jiang, S., Hillyer, C. & Du, L. Neutralizing antibodies against SARS-CoV-2 and other human coronaviruses. Trends Immunol. 41, 355–359 (2020).

19 Wajnberg, A. et al. Robust neutralizing antibodies to SARS-CoV-2 infection persist for months. Science 370, 1227–1230 (2020).

20 Yuan, M. et al. A highly conserved cryptic epitope in the receptor binding domains of SARS-CoV-2 and SARS-CoV. Science 368, 630–633 (2020).

21 Pinto, D. et al. Cross-neutralization of SARS-CoV-2 by a human monoclonal SARS-CoV antibody. Nature 583, 290–295 (2020).

22 Ju, B. et al. Human neutralizing antibodies elicited by SARS-CoV-2 infection. Nature 584, 115–119 (2020).

23 Cao, Y. et al. Potent neutralizing antibodies against SARS-CoV-2 identified by high-throughput single-cell sequencing of convalescent patients’ B cells. Cell 182, 73–84 (2020).

24 Rogers, T. F. et al. Isolation of potent SARS-CoV-2 neutralizing antibodies and protection from disease in a small animal model. Science 369, 956–963 (2020).

25 Brouwer, P. J. M. et al. Potent neutralizing antibodies from COVID-19 patients define multiple targets of vulnerability. Science 369, 643–650 (2020).

26 Wu, Y. et al. A noncompeting pair of human neutralizing antibodies block COVID-19 virus binding to its receptor ACE2. Science 368, 1274–1278 (2020).

27 Chi, X. et al. A neutralizing human antibody binds to the N-terminal domain of the Spike protein of SARS-CoV-2. Science 369, 650–655 (2020).

28 Seydoux, E. et al. Characterization of neutralizing antibodies from a SARS-CoV-2 infected individual. bioRxiv, doi:https://doi.org/10.1101/2020.05.12.091298 (2020).

29 Shi, R. et al. A human neutralizing antibody targets the receptor-binding site of SARS-CoV-2. Nature 584, 120–124 (2020).

30 Wan, J. et al. Human-IgG-neutralizing monoclonal antibodies block the SARS-CoV-2 infection. Cell Rep. 32, 107918 (2020).

31 Barnes, C. O. et al. Structures of human antibodies bound to SARS-CoV-2 spike reveal common epitopes and recurrent features of antibodies. Cell 182, 828–842 (2020).

32 Robbiani, D. F. et al. Convergent antibody responses to SARS-CoV-2 in convalescent individuals. Nature 584, 437–442 (2020).

33 Han, X. et al. A rapid and efficient screening system for neutralizing antibodies and its application for the discovery of potent neutralizing antibodies to SARS-CoV-2 S-RBD. bioRxiv, doi:https://doi.org/10.1101/2020.08.19.253369 (2020).

34 Kreye, J. et al. A therapeutic non-self-reactive SARS-CoV-2 antibody protects from lung pathology in a COVID-19 hamster model. Cell 183, 1058–1069 (2020).

35 Zost, S. J. et al. Rapid isolation and profiling of a diverse panel of human monoclonal antibodies targeting the SARS-CoV-2 spike protein. Nat. Med. 26, 1422–1427 (2020).

36 Liu, L. et al. Potent neutralizing antibodies against multiple epitopes on SARS-CoV-2 spike. Nature 584, 450–456 (2020).

37 Kreer, C. et al. Longitudinal isolation of potent near-germline SARS-CoV-2-neutralizing antibodies from COVID-19 patients. Cell 182, 843–854 (2020).

38 Hurlburt, N. K. et al. Structural basis for potent neutralization of SARS-CoV-2 and role of antibody affinity maturation. Nat. Commun. 11, 5413 (2020).

39 Noy-Porat, T. et al. A panel of human neutralizing mAbs targeting SARS-CoV-2 spike at multiple epitopes. Nat. Commun. 11, 4303 (2020).

40 Du, S. et al. Structurally resolved SARS-CoV-2 antibody shows high efficacy in severely infected hamsters and provides a potent cocktail pairing strategy. Cell 183, 1013–1023 (2020).

41 Hozumi, N. & Tonegawa, S. Evidence for somatic rearrangement of immunoglobulin genes coding for variable and constant regions. Proc. Natl. Acad. Sci. U. S. A. 73, 3628–3632 (1976).

42 Brack, C., Hirama, M., Lenhard-Schuller, R. & Tonegawa, S. A complete immunoglobulin gene is created by somatic recombination. Cell 15, 1–14 (1978).

43 Seidman, J. et al. Multiple related immunoglobulin variable-region genes identified by cloning and sequence analysis. Proc. Natl. Acad. Sci. U. S. A. 75, 3881–3885 (1978).

44 Bernard, O., Hozumi, N. & Tonegawa, S. Sequences of mouse immunoglobulin light chain genes before and after somatic changes. Cell 15, 1133–1144 (1978).

45 O’Brien, R., Brinster, R. & Storb, U. Somatic hypermutation of an immunoglobulin transgene in K transgenic mice. Nature 326, 405–409 (1987).

46 Nielsen, S. C. A. et al. Human B cell clonal expansion and convergent antibody responses to SARS-CoV-2. Cell Host Microbe 28, 516–525 (2020).

47 Yuan, M. et al. Structural basis of a shared antibody response to SARS-CoV-2. Science 369, 1119–1123 (2020).

48 Setliff, I. et al. Multi-donor longitudinal antibody repertoire sequencing reveals the existence of public antibody clonotypes in HIV-1 infection. Cell Host Microbe 23, 845–854 (2018).

49 Jackson, K. J. et al. Human responses to influenza vaccination show seroconversion signatures and convergent antibody rearrangements. Cell Host Microbe 16, 105–114 (2014).

50 Trück, J. et al. Identification of antigen-specific B cell receptor sequences using public repertoire analysis. J. Immunol. 194, 252–261 (2015).

51 Dunand, C. J. H. & Wilson, P. C. Restricted, canonical, stereotyped and convergent immunoglobulin responses. Philos. Trans. R. Soc. Lond., Ser. B: Biol. Sci. 370, 20140238 (2015).

52 Pieper, K. et al. Public antibodies to malaria antigens generated by two LAIR1 insertion modalities. Nature 548, 597–601 (2017).

53 Parameswaran, P. et al. Convergent antibody signatures in human dengue. Cell Host Microbe 13, 691–700 (2013).

54 Wu, N. C. et al. An alternative binding mode of IGHV3-53 antibodies to the SARS-CoV-2 receptor binding domain. Cell Rep. 33, 108274 (2020).

55 Barnes, C. O. et al. SARS-CoV-2 neutralizing antibody structures inform therapeutic strategies. Nature 588, 682–687 (2020).

56 Bertoglio, F. et al. A SARS-CoV-2 neutralizing antibody selected from COVID-19 patients by phage display is binding to the ACE2-RBD interface and is tolerant to known RBD mutations. bioRxiv, doi:https://doi.org/10.1101/2020.12.03.409318 (2020).

57 Clark, S. A. et al. Molecular basis for a germline-biased neutralizing antibody response to SARS-CoV-2. bioRxiv, doi:https://doi.org/10.1101/2020.11.13.381533 (2020).

58 Sinnokrot, M. O., Valeev, E. F. & Sherrill, C. D. Estimates of the ab initio limit for π-π interactions: The benzene dimer. J. Am. Chem. Soc. 124, 10887–10893 (2002).

59 Chelli, R., Gervasio, F. L., Procacci, P. & Schettino, V. Stacking and T-shape competition in aromatic-aromatic amino acid interactions. J. Am. Chem. Soc. 124, 6133–6143 (2002).

60 Kim, S. I. et al. Stereotypic neutralizing VH antibodies against SARS-CoV-2 spike protein receptor binding domain in COVID-19 patients and healthy individuals. Sci. Transl. Med., doi:https://doi.org/10.1126/scitranslmed.abd6990 (2021).

61 Banach, B. B. et al. Paired heavy and light chain signatures contribute to potent SARS-CoV-2 neutralization in public antibody responses. bioRxiv, doi:https://doi.org/10.1101/2020.12.31.424987 (2021).

62 Gaebler, C. et al. Evolution of antibody immunity to SARS-CoV-2. Nature, doi:https://doi.org/10.1038/s41586-021-03207-w (2021).

63 Tegally, H. et al. Emergence and rapid spread of a new severe acute respiratory syndrome-related coronavirus 2 (SARS-CoV-2) lineage with multiple spike mutations in South Africa. medRxiv, doi:https://doi.org/10.1101/2020.12.21.20248640 (2020).

64 Voloch, C. M. et al. Genomic characterization of a novel SARS-CoV-2 lineage from Rio de Janeiro, Brazil. medRxiv, doi:https://doi.org/10.1101/2020.12.23.20248598 (2020).

65 Weisblum, Y. et al. Escape from neutralizing antibodies by SARS-CoV-2 spike protein variants. eLife 9, e61312(2020).

66 Greaney, A. J. et al. Complete mapping of mutations to the SARS-CoV-2 spike receptor-binding domain that escape antibody recognition. Cell Host Microbe 29, 44–57 (2021).

67 Andreano, E. et al. SARS-CoV-2 escape in vitro from a highly neutralizing COVID-19 convalescent plasma. bioRxiv, doi:https://doi.org/10.1101/2020.12.28.424451 (2020).

68 Buss, L. F. et al. Three-quarters attack rate of SARS-CoV-2 in the Brazilian Amazon during a largely unmitigated epidemic. Science 371, 288–292 (2021).

69 Greaney, A. J. et al. Comprehensive mapping of mutations to the SARS-CoV-2 receptor-binding domain that affect recognition by polyclonal human serum antibodies. bioRxiv, doi:https://doi.org/10.1101/2020.12.31.425021 (2021).

70 Wang, Z. et al. mRNA vaccine-elicited antibodies to SARS-CoV-2 and circulating variants. bioRxiv, 2021.2001.2015.426911, doi:10.1101/2021.01.15.426911 (2021).

71 Wibmer, C. K. et al. SARS-CoV-2 501Y.V2 escapes neutralization by South African COVID-19 donor plasma. bioRxiv, 2021.2001.2018.427166, doi:10.1101/2021.01.18.427166 (2021).

72 Ye, J., Ma, N., Madden, T. L. & Ostell, J. M. IgBLAST: an immunoglobulin variable domain sequence analysis tool. Nucleic Acids Res. 41, W34–W40 (2013).

73 Crooks, G. E., Hon, G., Chandonia, J.-M. & Brenner, S. E. WebLogo: a sequence logo generator. Genome Res. 14, 1188–1190 (2004).

74 Wu, N. C. et al. In vitro evolution of an influenza broadly neutralizing antibody is modulated by hemagglutinin receptor specificity. Nat. Commun. 8, 15371 (2017).

75 Otwinowski, Z. & Minor, W. Processing of X-ray diffraction data collected in oscillation mode. Methods Enzymol. 276, 307–326 (1997).

76 McCoy, A. J. et al. Phaser crystallographic software. J. Appl. Crystallogr. 40, 658–674 (2007).

77 Schritt, D. et al. Repertoire Builder: high-throughput structural modeling of B and T cell receptors. Mol. Syst. Des. Eng. 4, 761–768, doi:10.1039/C9ME00020H (2019).

78 Emsley, P., Lohkamp, B., Scott, W. G. & Cowtan, K. Features and development of Coot. Acta Crystallogr. D Biol. Crystallogr. 66, 486–501 (2010).

79 Adams, P. D. et al. PHENIX: a comprehensive Python-based system for macromolecular structure solution. Acta Crystallogr. D Biol. Crystallogr. 66, 213–221 (2010).

80 Chao, G. et al. Isolating and engineering human antibodies using yeast surface display. Nat. Protoc. 1, 755 (2006).

81 Benatuil, L., Perez, J. M., Belk, J. & Hsieh, C.-M. An improved yeast transformation method for the generation of very large human antibody libraries. Protein Eng. Des. Sel. 23, 155–159 (2010).

82 Zhang, J., Kobert, K., Flouri, T. & Stamatakis, A. PEAR: a fast and accurate Illumina Paired-End reAd mergeR. Bioinformatics 30, 614–620 (2014).

83 Kovaltsuk, A. et al. Observed antibody space: A resource for data mining next-generation sequencing of antibody repertoires. J. Immunol. 201, 2502–2509 (2018).

