## Supplementary Information for "Sequence signatures of two IGHV3-53/3-66 public clonotypes to SARS-CoV-2 receptor binding domain"

### Supplementary Figures

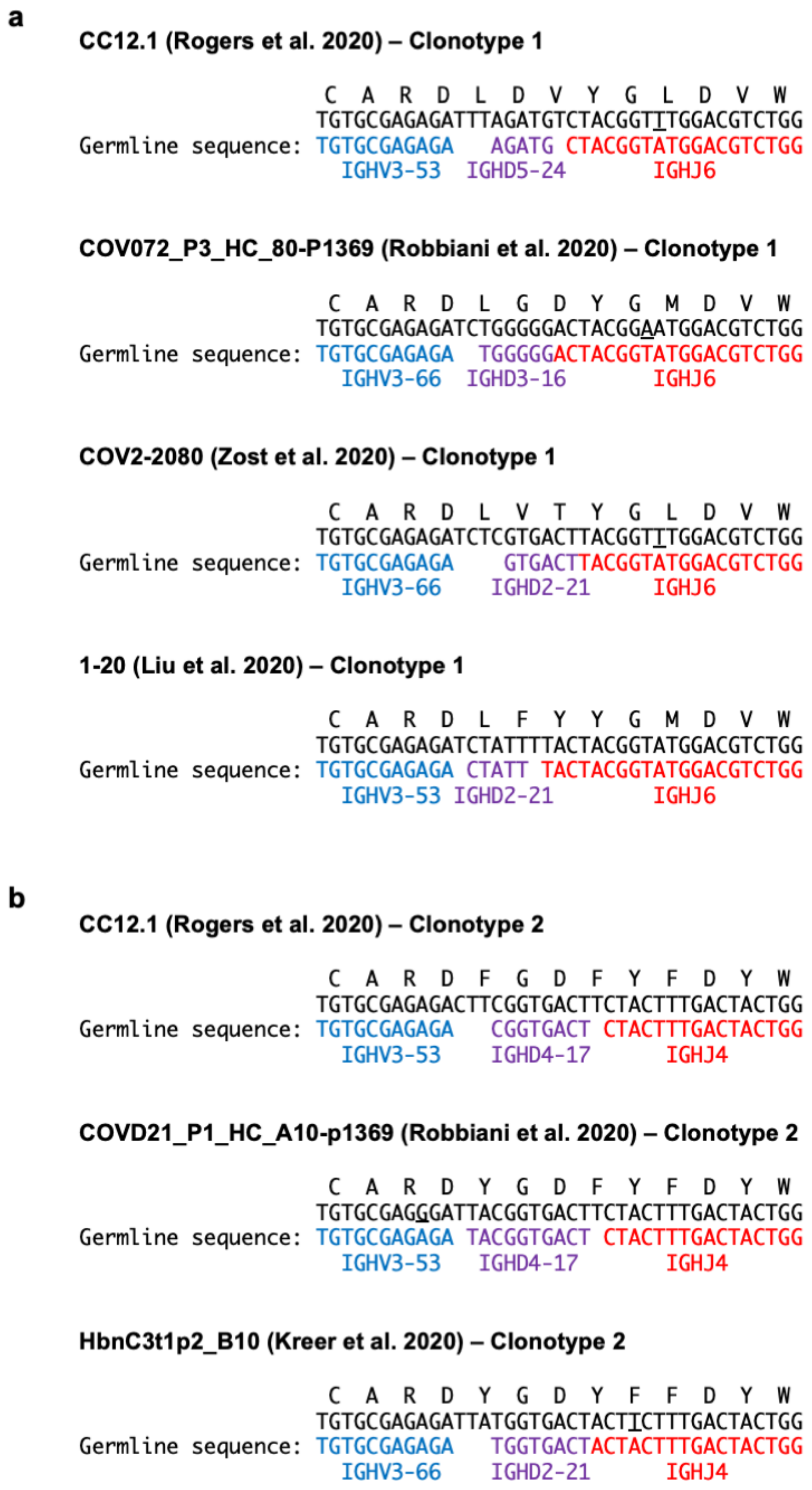

**Supplementary Figure 1. Putative germline sequences for CDR H3 in representative antibodies from clonotypes 1 and 2.** Amino acid and nucleotide sequences of the V-D-J junction of **(a)** four clonotype 1 antibodies from different studies, and **(b)** three clonotype 2 antibodies from different studies, are shown. Putative germline sequences and segments are indicated. Somatically mutated nucleotides are underlined. Intervening spaces at the V-D and D-J junctions are N-nucleotide additions.

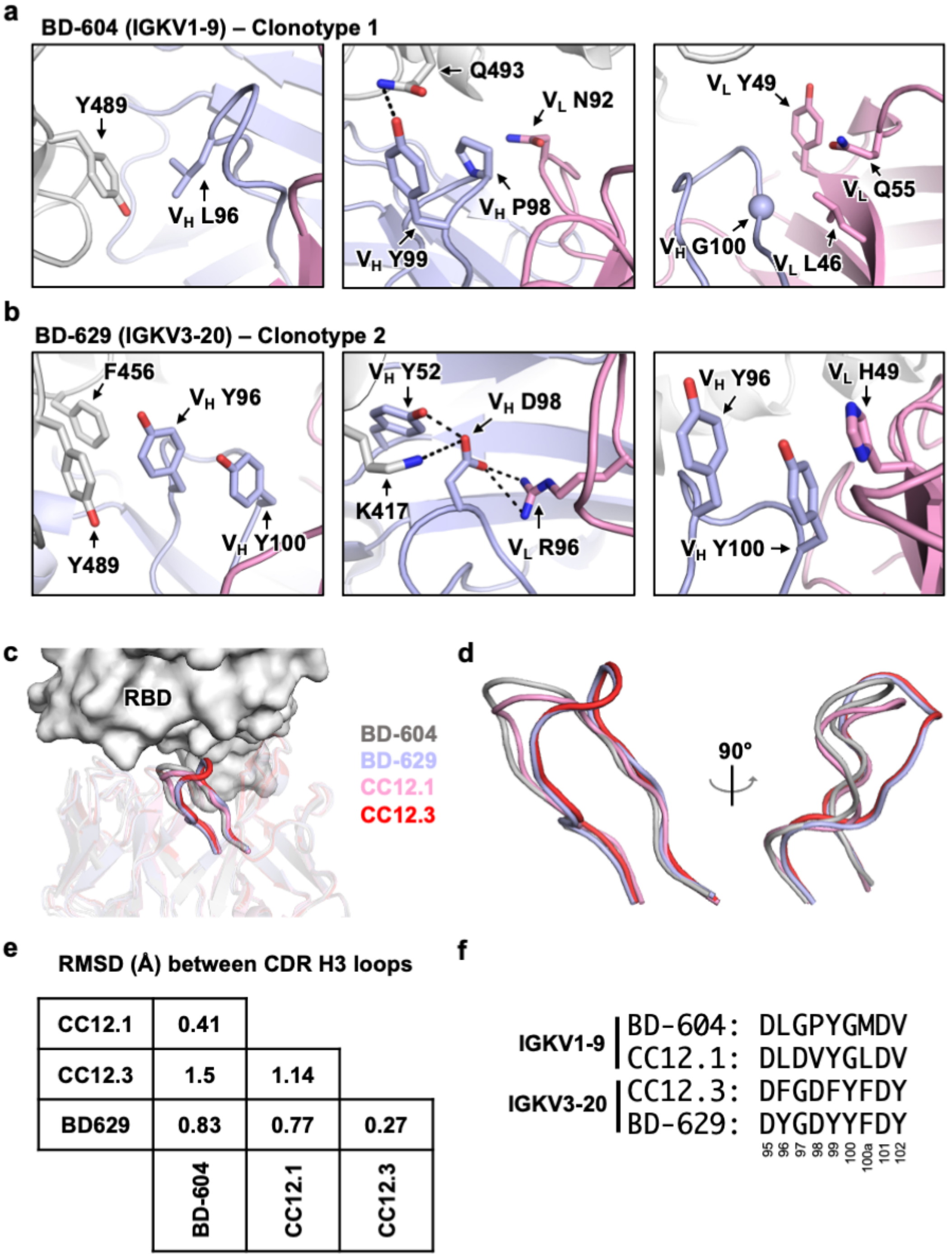


**Supplementary Figure 2. Interactions between CDR H3 and the light chain of two clonotypes. (a)** Interactions of L96, V98 and G100 (Kabat numbering) in CDR H3 of BD-604 (PDB 7CH4) with the IGKV1-9 light chain, and SARS-CoV-2 RBD. **(b)** Interactions of Y96, D98 and Y100 (Kabat numbering) in CDR H3 of BD-629 (PDB 7CH5) with the IGKV3-20 light chain, and SARS-CoV-2 RBD. Gray: RBD; light blue: heavy chain; pink: light chain. **(c-d)** Structural alignment of four antibodies highlighting the structural similarities and differences of the CDR H3 loop. Gray: BD-604; light blue: BD-629; pink: CC12.1; red: CC12.3. **(c)** An overall view with RBD as white surface and antibodies as cartoon representations. The antibodies are semi-transparent except for the CDR H3 loops. **(d)** Zoomed-in views of the CDR H3 loops. **(e)** Root-mean-square deviation (RMSD) are shown for the atomic positions of the CDR H3 loops from different antibodies. **(f)** Sequences of CDR H3 loops from different antibodies are shown. Amino acids are labeled according to Kabat numbering.

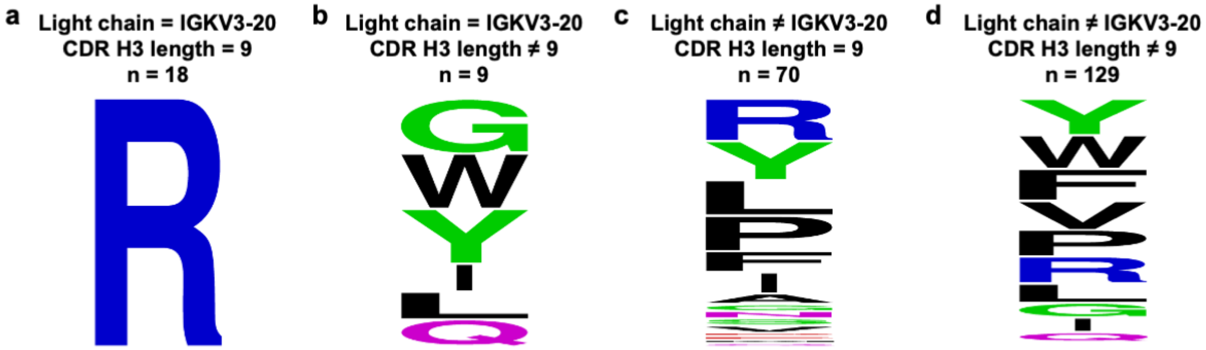


**Supplementary Figure 3. Sequence logos of residue 96 in CDR L3 of IGHV3-53/3-66 RBD antibodies. (a)** IGKV3-20 as light chain and CDR H3 length of 9 (n = 18), **(b)** IGKV3-20 as light chain and CDR H3 length not 9 (n = 9), **(c)** non-IGKV3-20 as light chain and CDR H3 length of 9 (n = 70), and **(d)** non-IGKV3-20 as light chain and CDR H3 length not equal to 9 (n = 129).

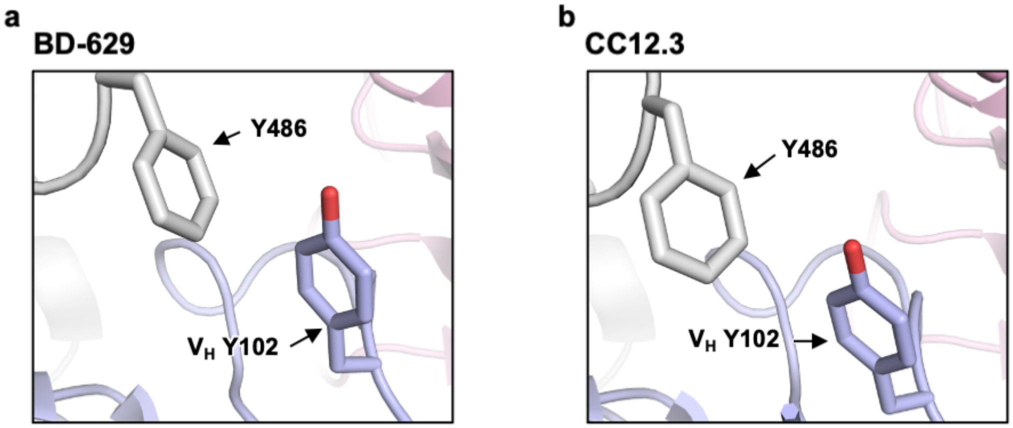


**Supplementary Figure 4. Interactions between Y102 of CDR H3 loop of class 2 antibodies and the RBD of the S protein. (a)** Interaction of Y102 (Kabat numbering) in CDR H3 of BD-629 (PDB 7CH5) with the RBD of the S protein. **(b)** Interaction of Y102 (Kabat numbering) in CDR H3 of CC12.3 (PDB 6XC4) with the RBD of the S protein.

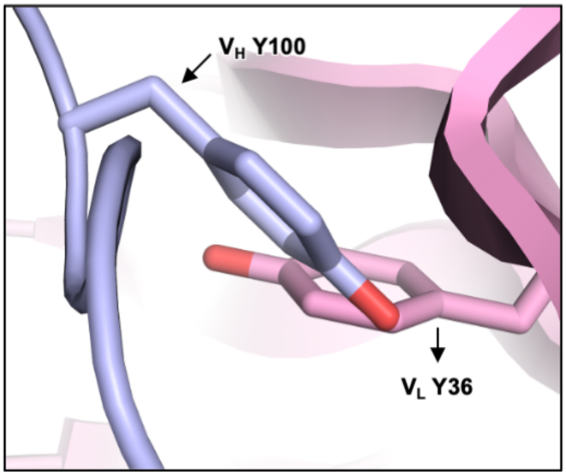


**Supplementary Figure 5. V$_H$ Y100 would clash with the light chain.** Rosetta modeling[1] of G100 of CC12.1 (PDB 6XC2), a clonotype 1 antibody, to Y100 with a fixed backbone indicates a clash of V$_H$ Y100 with the V$_L$ Y36.

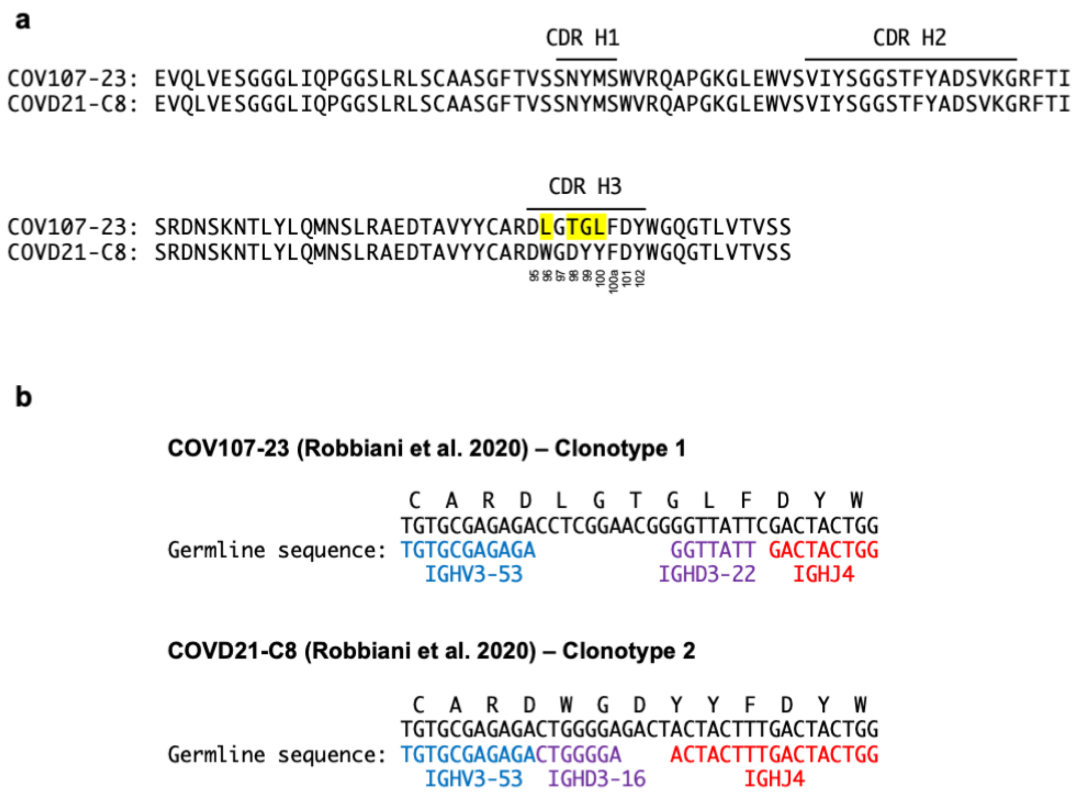


**Supplementary Figure 6. Putative germline sequences for CDR H3 in representative antibodies from clonotypes 1 and 2. (a)** The heavy chains of COV107-23 (clonotype 1) and COVD21-C8 (clonotype 2) RBD antibodies from Robbiani et al.[2] are aligned as shown. CDRs are annotated based on Kabat numbering. **(b)** Amino acid and nucleotide sequences of the V-D-J junction of COV107-23 and COVD21-C8 are shown. Putative germline sequences and segments are indicated. COV107-23 and COVD21-C8 are named as COV107_Plate2_HC_23-P1369 and COVD21_P1_HC_C8-p1369, respectively, in Supplementary Table 1 of Robbiani et al.[2]

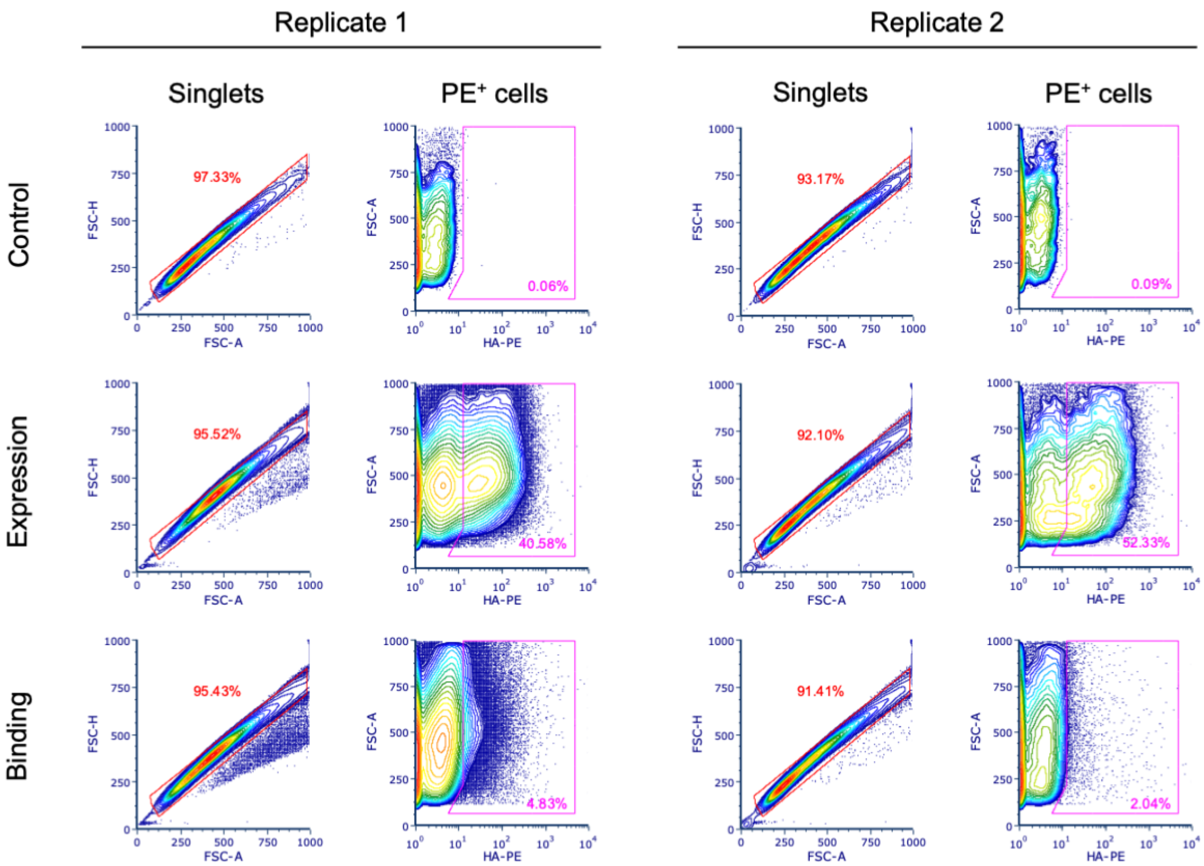


**Supplementary Figure 7. Density plots of fluorescence-activated cell sorting of yeast cells.** Cells from the yeast display library were sorted based on signal of PE. PE-positive singlets were collected. In the first replicate, the singlet gate was unrestricted, while in the second replicate, the singlet gate was restricted to collect 100,000 cells.

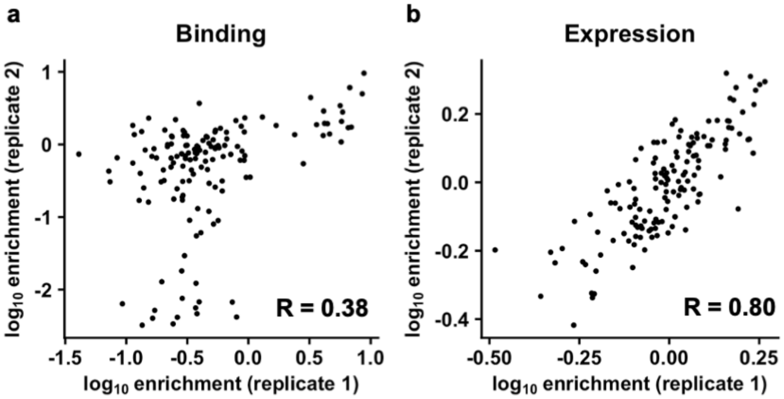


**Supplementary Figure 8. Correlation between biological replicates of the yeast display selection experiment for 143 CDR H3 variants in the B38 antibody. (a)** Correlation of $\log_{10}$ enrichment in binding between biological replicates is shown. **(b)** Correlation of $\log_{10}$ enrichment in expression between biological replicates is shown. The Pearson correlation (R) is indicated.

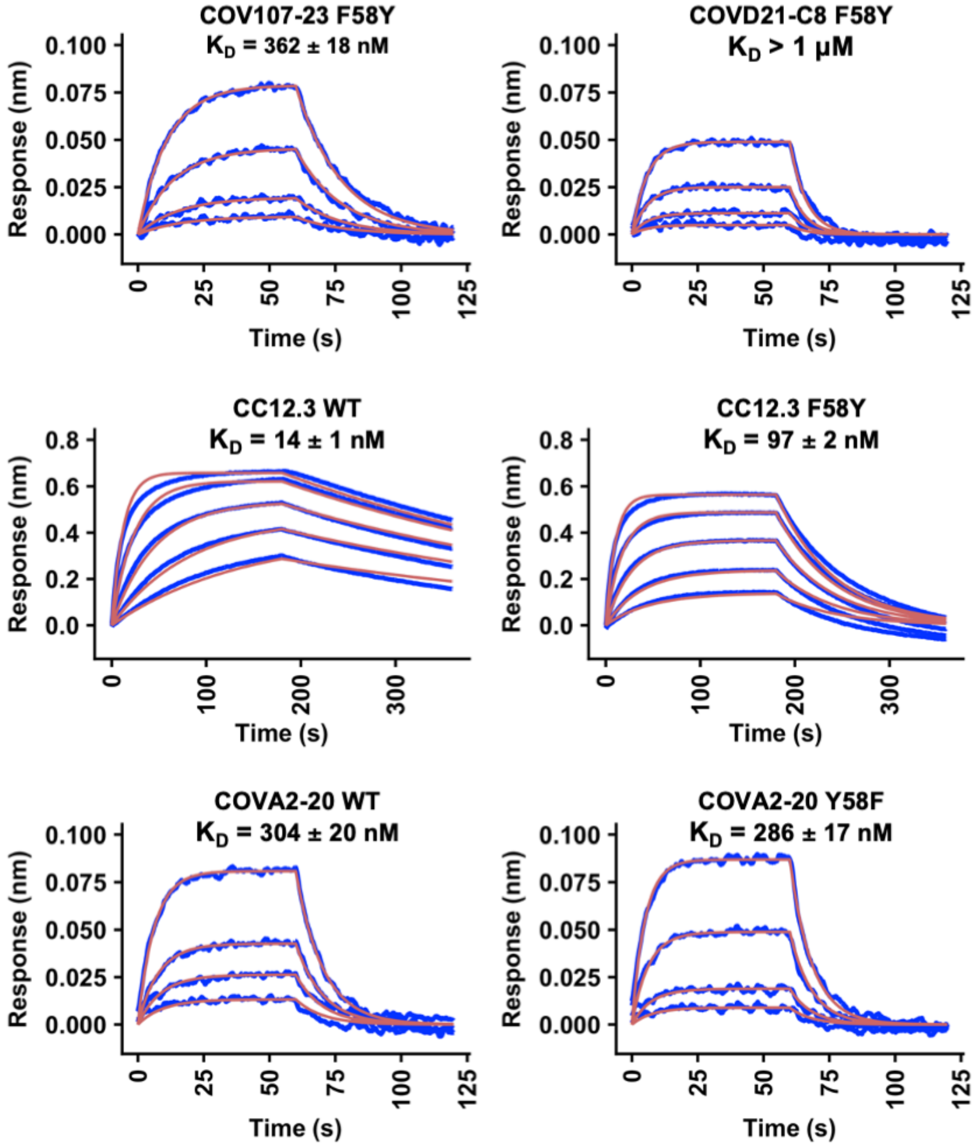


**Supplementary Figure 9. Sensorgrams for binding of Fabs to SARS-CoV-2 RBD.** Binding kinetics of different Fabs against recombinant SARS-CoV-2 RBD were measured by biolayer interferometry (BLI). Y-axis represents the response. Blue lines represent the response curve and red lines represent a 1:1 binding model. Binding kinetics were measured for four to five Fab concentrations.

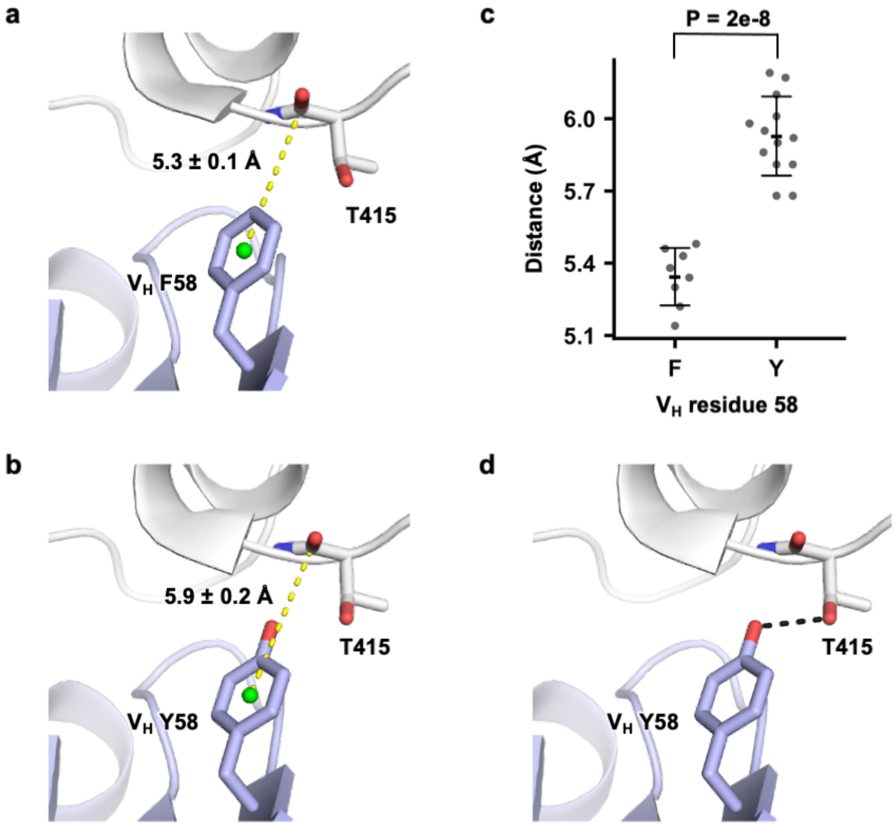


**Supplementary Figure 10. Structural analysis of the Y58F mutation. (a)** The distance between the centroid of the aromatic ring of $V_H$ F58 and the backbone carbon atom of RBD T415 was measured for seven IGHV3-53/66 RBD antibodies with Y58F mutation, namely P4A1 (PDB 7CJF), CC12.1 (PDB 6XC3), CC12.3 (PDB 6XC4), C102 (PDB 7K8M), BD-604 (PDB 7CH4), COVA2-04 (PDB 7JMO), and CB6 (PDB 7C01). (n.b., there are two copies of CC12.3-RBD complex in PDB 6XC4 and both copies were analyzed). The mean and standard deviation are indicated. The centroid of the aromatic ring is represented by the green sphere. **(b)** Same as panel **a**, except for nine IGHV3-53/3-66 RBD antibodies without a Y58F mutation, namely CV30 (PDB 6XE1), STE90-C11 (PDB 7B3O), B38 (PDB 7BZ5), P2C-1F11 (PDB 7CDI), C1A-B3 (PDB 7KFW), BD-629 (PDB 7CH5), C1A-F10 (PDB 7KFY), C1A-B12 (PDB 7KFV), and C1A-C2 (PDB 7KFX). (n.b., there are three copies of C1A-B3-RBD complex in PDB 7KFW and three copies of C1A-B12-RBD complex in PDB 7KFV, all of which were analyzed). **(c)** The distance data reported in panels **a** and **b** are shown. **(d)** The hydroxyl group of $V_H$ Y58 can form a H-bonds (black dashed line) with RBD T415. The hydrogen bond is identified by PISA (https://www.ebi.ac.uk/pdbe/pisa/)[3].

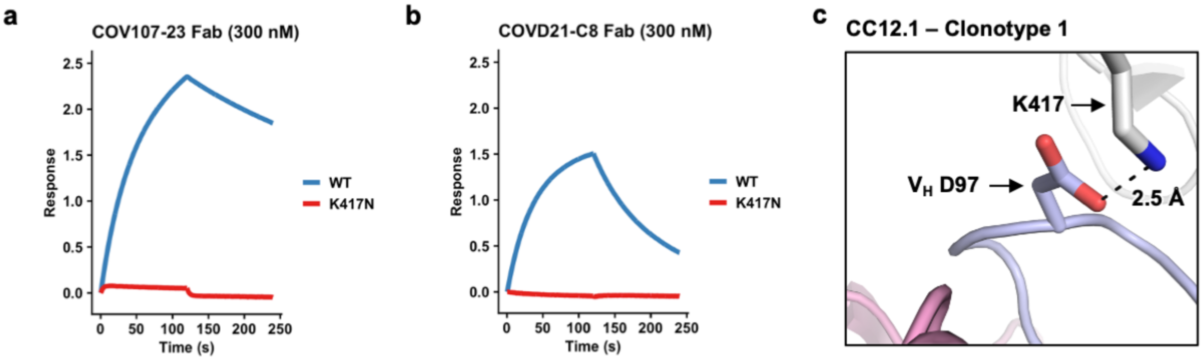


**Supplementary Figure 11. Clonotypes 1 and 2 strongly bind to wild-type SARS-CoV-2 RBD but not to SARS-CoV-2 K417N RBD mutant.** Binding kinetics of **(a)** COV107-23 (clonotype 1) and **(b)** COVD21-C8 (clonotype 2) Fabs to recombinant SARS-CoV-2 RBD were measured by biolayer interferometry. Y-axis represents the response. Blue and red lines represent the response curve for binding of Fabs to wild-type (WT) and K417N RBD mutant, respectively. **(c)** Electrostatic interaction between $V_H$ D97 of CC12.1 (PDB 6XC2), a clonotype 1 antibody, with K417 RBD.

### Supplementary Tables

**Supplementary Table 1. IGHV3-53/3-66 RBD antibodies.** List of 214 published IGHV3-53/3-66 RBD antibodies obtained from convalescent patients infected with SARS-CoV-2.

**Supplementary Table 2. X-ray data collection and refinement statistics**

| **Data collection** | | |
|---|---|---|
| | COVA107-23 WT | COVA107-23 HC + COVD21-C8 LC |
| Beamline | SSRL 12-1 | SSRL 12-1 |
| Wavelength (Å) | 0.97946 | 0.97946 |
| Space group | P 2 | P 1 |
| Unit cell parameters | | |
| a, b, c (Å) | 79.4, 74.8, 172.7 | 73.3, 78.4, 91.6 |
| α, β, γ (°) | 90, 99.4, 90 | 100.1, 112.1, 92.8 |
| Resolution (Å)[a] | 50.0–1.98 (2.01–1.98) | 50.0–3.30 (3.40–3.30) |
| Unique reflections[a] | 130,795 (12,808) | 24,557 (2,165) |
| Redundancy[a] | 3.5 (2.9) | 1.7 (1.6) |
| Completeness (%)[a] | 94.3 (94.5) | 82.3 (70.7) |
| $\langle I/\sigma_I \rangle$[a] | 13.8 (1.0) | 2.7 (1.0) |
| $R_{sym}$[b] (%)[a] | 14.0 (>100) | 29.9 (70.9) |
| $R_{pim}$[b] (%)[a] | 5.7 (64.0) | 18.8 (46.4) |
| $CC_{1/2}$[c] (%)[a] | 99.7 (66.3) | 93.4 (72.7) |
| **Refinement statistics** | | |
| Resolution (Å) | 30.7–1.98 | 35.9–3.30 |
| Reflections (work) | 130,416 | 24,493 |
| Reflections (test) | 6,632 | 1,186 |
| $R_{cryst}$[d] / $R_{free}$[e] (%) | 21.4/26.3 | 29.8/31.9 |
| No. of atoms | 14,029 | 12,881 |
| Macromolecules | 12,888 | 12,881 |
| Solvent | 1,137 | 0 |
| Average *B*-value ($Å^2$) | 38 | 59 |
| Macromolecules | 38 | 59 |
| Solvent | 41 | N/A |
| Wilson *B*-value ($Å^2$) | 30 | 54 |
| **RMSD from ideal geometry** | | |
| Bond length (Å) | 0.012 | 0.014 |
| Bond angle (°) | 1.29 | 0.74 |
| **Ramachandran statistics (%)** | | |
| Favored | 95.5 | 95.2 |
| Outliers | 0.06 | 0.00 |
| **PDB code** | Pending | Pending |

[a] Numbers in parentheses refer to the highest resolution shell.

[b] $R_{sym} = \Sigma_{hkl} \Sigma_i | I_{hkl,i} - \langle I_{hkl} \rangle | / \Sigma_{hkl} \Sigma_i I_{hkl,i}$ and $R_{pim} = \Sigma_{hkl} (1/(n-1))^{1/2} \Sigma_i | I_{hkl,i} - \langle I_{hkl} \rangle | / \Sigma_{hkl} \Sigma_i I_{hkl,i}$, where $I_{hkl,i}$ is the scaled intensity of the $i^{th}$ measurement of reflection h, k, l, $\langle I_{hkl} \rangle$ is the average intensity for that reflection, and $n$ is the redundancy.

[c] $CC_{1/2}$ = Pearson correlation coefficient between two random half datasets.

[d] $R_{cryst} = \Sigma_{hkl} | F_o - F_c | / \Sigma_{hkl} | F_o | \times 100$, where $F_o$ and $F_c$ are the observed and calculated structure factors, respectively.

[e] $R_{free}$ was calculated as for $R_{cryst}$, but on a test set comprising 5% of the data excluded from refinement.

**Supplementary Table 3. Oligonucleotide sequences encoding CDR H3 loops.** List of 143 oligonucleotides encoding CDR H3 loops used in generation of a yeast antibody display library.

**Supplementary Table 4. Read count and enrichment level of each CDR H3 variant in the yeast display screen.**

### Supplementary References


1. Kellogg, E. H., Leaver-Fay, A. & Baker, D. Role of conformational sampling in computing mutation-induced changes in protein structure and stability. *Proteins: Struct. Funct. Bioinform.* **79**, 830-838 (2011).

2. Robbiani, D. F. *et al.* Convergent antibody responses to SARS-CoV-2 in convalescent individuals. *Nature* **584**, 437-442 (2020).

3. Krissinel, E. & Henrick, K. Inference of macromolecular assemblies from crystalline state. *J. Mol. Biol.* **372**, 774-797 (2007).
